# An siRNA-guided Argonaute protein directs RNA Polymerase V to initiate DNA methylation

**DOI:** 10.1101/2021.08.19.457027

**Authors:** Meredith J. Sigman, Kaushik Panda, Rachel Kirchner, Lauren L. McLain, Hayden Payne, John Reddy Peasari, Aman Y. Husbands, R. Keith Slotkin, Andrea D. McCue

**Affiliations:** Department of Molecular Genetics, The Ohio State University, Columbus, Ohio, 43210, USA; Donald Danforth Plant Science Center, St. Louis, Missouri, 63132, USA; Bioinformatics and Computational Biology Program, Saint Louis University, St. Louis, Missouri, 63108, USA; Division of Biological Sciences, University of Missouri, Columbia, Missouri, 65211, USA

**Keywords:** RNA-directed DNA methylation (RdDM), small interfering RNA (siRNA), RNA Polymerase V, Argonaute, epigenetics, DNA methylation

## Abstract

In mammals and plants, cytosine DNA methylation is essential for the epigenetic repression of transposable elements and foreign DNA. In plants, DNA methylation is guided by small interfering RNAs (siRNAs) in a self-reinforcing cycle termed RNA-directed DNA methylation (RdDM). RdDM requires the specialized RNA Polymerase V (Pol V), and the key unanswered question is how Pol V is first recruited to new target sites without preexisting DNA methylation. We find that Pol V follows and is dependent upon the recruitment of an AGO4-clade ARGONAUTE protein, and any siRNA can guide the ARGONAUTE protein to the new target locus independent of preexisting DNA methylation. These findings reject long-standing models of RdDM initiation and instead demonstrate that siRNA-guided ARGONAUTE targeting is necessary, sufficient and first to target Pol V recruitment and trigger the cycle of RdDM at a transcribed target locus, thereby establishing epigenetic silencing.

---

Chromatin marks segregate genomes into expressed domains and regions that remain transcriptionally silenced. In mammals and plants, DNA methylation provides information such as which regions are transposable elements (TEs) and should not be expressed (reviewed in ^1^), while integrated transgenes are often subject to this regulation as well^2^. Most studies have focused on how DNA methylation is epigenetically maintained, resulting in heritable transcriptional repression. However, how DNA methylation is initially established at individual loci is less understood.

In both plants and mammals, DNA methylation is targeted via the action of small RNAs (piRNAs in mammals)^3,4^. Specifically in plants, small interfering RNAs (siRNAs) are produced from TEs, viruses and transgenes, targeting them for RNA-directed DNA methylation (RdDM)(reviewed in ^5^). RdDM is a feed-forward cycle that reinforces DNA methylation and results in epigenetic transcriptional repression. The mechanism of RdDM is split between an upstream siRNA-generating phase and a downstream chromatin-linked phase. In the upstream phase, siRNAs are generated from either RNA Polymerase IV (Pol IV)- or Pol II-derived transcripts. In the downstream phase, these siRNAs are incorporated into one of the closely related ARGONAUTE proteins AGO4, AGO6 or AGO9^6^. Base complementarity between the siRNA and a chromatin-linked nascent transcript results in recruitment of the *de novo* DNA methyltransferases DRM1 and DRM2^7^. The nascent transcript is produced by RNA Polymerase V (Pol V), which provides the scaffold for AGO/siRNA complex interaction^8,9^. Pol IV and Pol V are derived from Pol II, as subunits of these holoenzymes duplicated early in plant evolution and subfunctionalized into their respective roles in siRNA biogenesis (Pol IV) and scaffolding RNA production (Pol V)^10^.

Pol V is continually recruited to RdDM loci through its interaction with the proteins SUVH2 and SUVH9, which bind existing cytosine DNA methylation^11,12^. However, how Pol V is recruited to new unmethylated DNA to first trigger RdDM and establish chromatin marks is not understood. One popular model in the literature proposes that, in the absence of Pol V, a Pol II-derived transcript acts as the scaffold and can set the initial round of DNA methylation^13,14^. A second model suggests that Pol V ubiquitously surveys the genome at low levels, and the first round of RdDM occurs upon addition of siRNAs. This ‘Pol V surveillance’ model is supported by a recent publication demonstrating that Pol V is present at a second set of loci that do not undergo RdDM because they lack siRNAs^15^. In both models, after the first round of methylation, Pol V would then be recruited through the activity of SUVH2/SUVH9, leading to the positive feed-forward loop of RdDM. Neither of the two models have been examined in the context of true ‘*de novo*’ silencing as in ^16^, and this leaves a formidable gap in our understanding of how the first round of DNA methylation is established.

Since RdDM is a self-reinforcing cycle, it is impossible to study the first round for endogenous regions of the genome that are already engaged. To address this, we developed an approach that interrogates *de novo* DNA methylation of newly transformed transgene DNA, so that no preexisting chromatin marks are guiding RdDM. We used this system to address a critical question in plant epigenetics: how is Pol V first recruited to new unmethylated sites for the first round of RdDM? Our findings demonstrate that, contrary to previous models, AGO4-clade proteins precede Pol V recruitment to new targets of RdDM. This mechanism provides the missing link between unmethylated DNA and the initiation of chromatin modification towards epigenetic silencing.

## Results

### A transgenic system to investigate the first round of DNA methylation

The expression of TE-derived sequences reproducibly triggers RdDM^17,18^. To study the initiation of DNA methylation, we recreated the ‘35S:EVD’ transgene consisting of a broadly expressed promoter driving the full-length coding sequence of the Arabidopsis *Evadé* TE^19^. This transgene was stably integrated into wild-type Columbia (wt Col) plant genomes, and transgene silencing in the first-generation (T1) transformants was assayed for all experiments. To determine if our TE transgene triggers RdDM, we utilized bisulfite amplicon sequencing (BSAS)^20^, a technique with high sequencing depth. In a side-by-side comparison, the average methylation remains the same for BSAS, Sanger sequencing and whole-genome sequencing techniques at target loci after conversion with sodium bisulfite, but the resolution of the data with BSAS is superior (∼3800 average coverage)(Supplemental Figure 1). When BSAS and small RNA sequencing were applied to T1 35:EVD plants, we found that siRNAs are generated from the transgene (Supplemental Figure 2), which results in high levels of *de novo* DNA methylation (Figure 1A and Supplemental Figure 2). Furthermore, although the coding sequence in 35S:EVD is an exact copy of the *Evadé* element EVD5 from the Arabidopsis genome (At5TE20395), transgene methylation is not influenced in *trans* by siRNAs from the endogenous EVD elements (Supplemental Figure 2).

**Figure 1.**
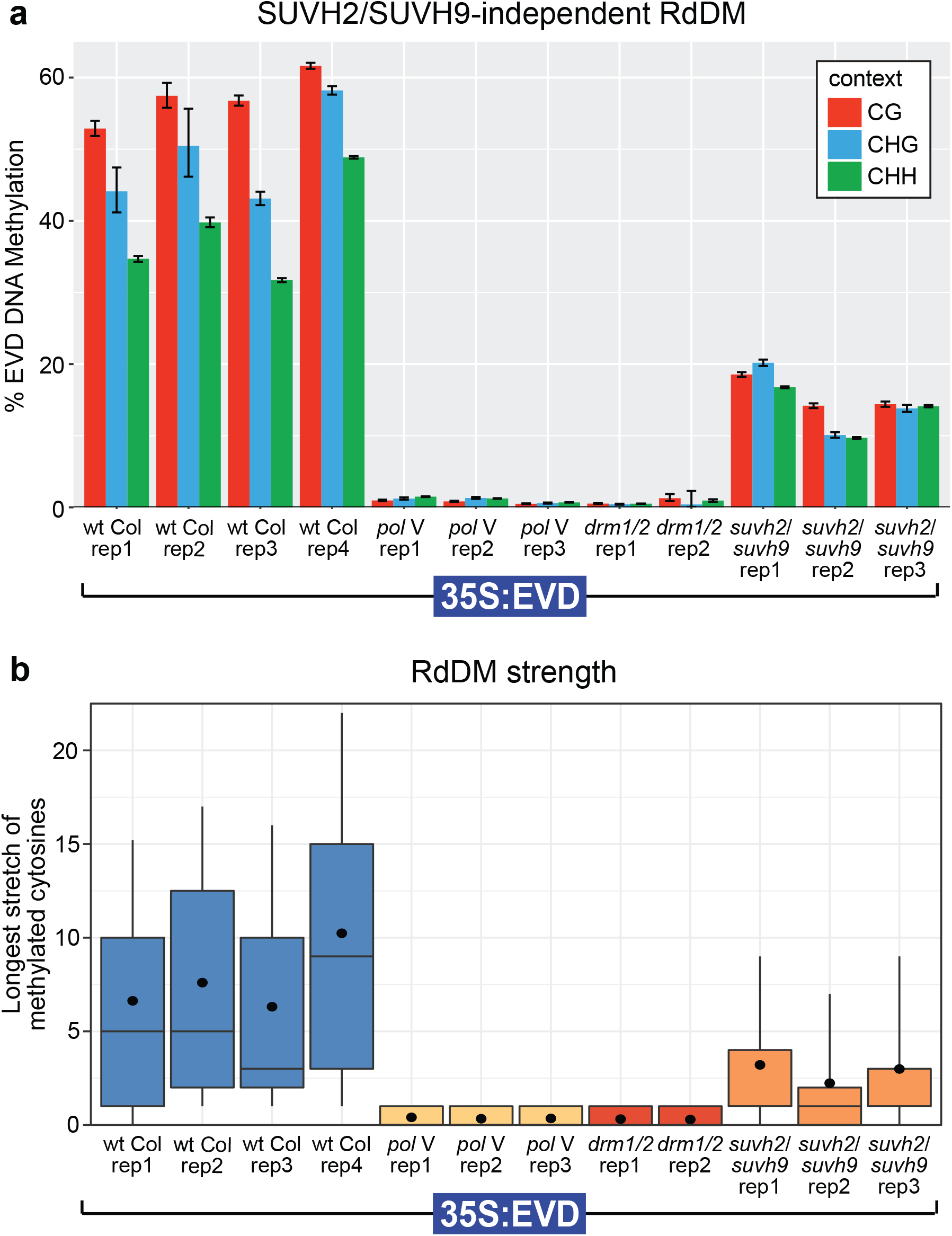
SUVH2/SUVH9-independent function of Pol V. A. Bisulfite Amplicon Sequencing (BSAS) of the 35S:EVD transgene in independent biological replicates of T1 plants. Error bars represent 95% Wilson confidence intervals. H = A, C or T. B. RdDM strength plot of DNA methylation. RdDM strength was measured by calculating the length of stretches of consecutively methylated cytosines along individual sequence reads. Box plots represent 25th and 75th percentile values with whiskers at the 10th and 90th percentile, the median is represented by a line, and the mean is denoted by a filled circle.

### SUVH2/SUVH9-independent Pol V function

Pol II has been proposed to substitute for Pol V during the first round of RdDM. To test this, we interrogated the methylation of T1 35S:EVD in *pol V* mutants (*nrpE1* subunit mutation). We found that Pol V is necessary to trigger RdDM, as plants lacking Pol V have only background levels of methylation similar to plants lacking *de novo* DNA methyltransferase activity (in the *drm1/2* double mutant), and similar to the non-conversion rate during sodium bisulfite treatment (Figure 1A)(non-conversion rates calculated in Supplemental Figure 1). BSAS affords increased data resolution^20^, which reveals that the low level of DNA methylation in *pol V* mutants exists as sporadic unconverted cytosines (stochastic), rather than as consecutive runs of methylated cytosines (indicative of DRM1/DRM2 activity in RdDM)(Supplemental Figure 1). We assayed RdDM strength by quantifying these strings of consecutive methylated bases (see Methods and controls in Supplemental Figure 1), which takes advantage of the large number of reads generated by BSAS to individually score each read, rather than averaging the data as in other methylation analysis methods. Using this improved methodology, we found no evidence of RdDM activity in *pol V* mutants (Figure 1B). Our data demonstrates that, contrary to the Pol II substitution model, Pol V is essential to initiate the first round of DNA methylation.

The only known mechanism for directly recruiting Pol V is through the DNA methyl-binding proteins SUVH2/SUVH9^11,12^. We found that the DNA methylation of 35S:EVD in plants lacking both SUVH2 and SUVH9 does not phenocopy the total loss of RdDM in *pol V* or *drm1/2* mutants (Figure 1A). Rather, *suvh2/suvh9* double mutants have an intermediate level of DNA methylation and RdDM strength (Figure 1A-B), suggesting that these proteins function to amplify DNA methylation levels rather than trigger RdDM at new locations. This demonstrates that there is a mechanism other than SUVH2/SUVH9 for the first round function of Pol V at an unmethylated target locus.

### AGO4-clade proteins are required for the first round of RdDM

To determine what other pathway components are essential for the first round of RdDM, we transformed 35S:EVD into a series of known DNA methylation, RdDM and RNAi mutants. As expected, 35S:EVD methylation and siRNA production are not dependent on DNA methyltransferases that propagate DNA methylation during DNA replication (maintenance methyltransferases)(Supplemental Figure 3). Rather, its methylation is dependent on at least one of the closely related AGO4-clade proteins (AGO4, AGO6 or AGO9 in the *ago4/6/9* triple mutant)(Figure 2A, Supplemental Figure 4). Because AGO2 has a published role in RdDM^21^, we additionally tested *ago2* mutants and found only partial reductions in 35S:EVD 21-22 nt siRNAs, DNA methylation and RdDM strength, suggesting it has a secondary role during the initiation of DNA methylation (Supplemental Figure 5). Furthermore, we used immunoprecipitation followed by small RNA sequencing and found that 35S:EVD-derived siRNAs are enriched in AGO4 protein complexes (Figure 2C, controls in Supplemental Figure 6), demonstrating the direct role of AGO4 in the RdDM of 35S:EVD.

**Figure 2.**
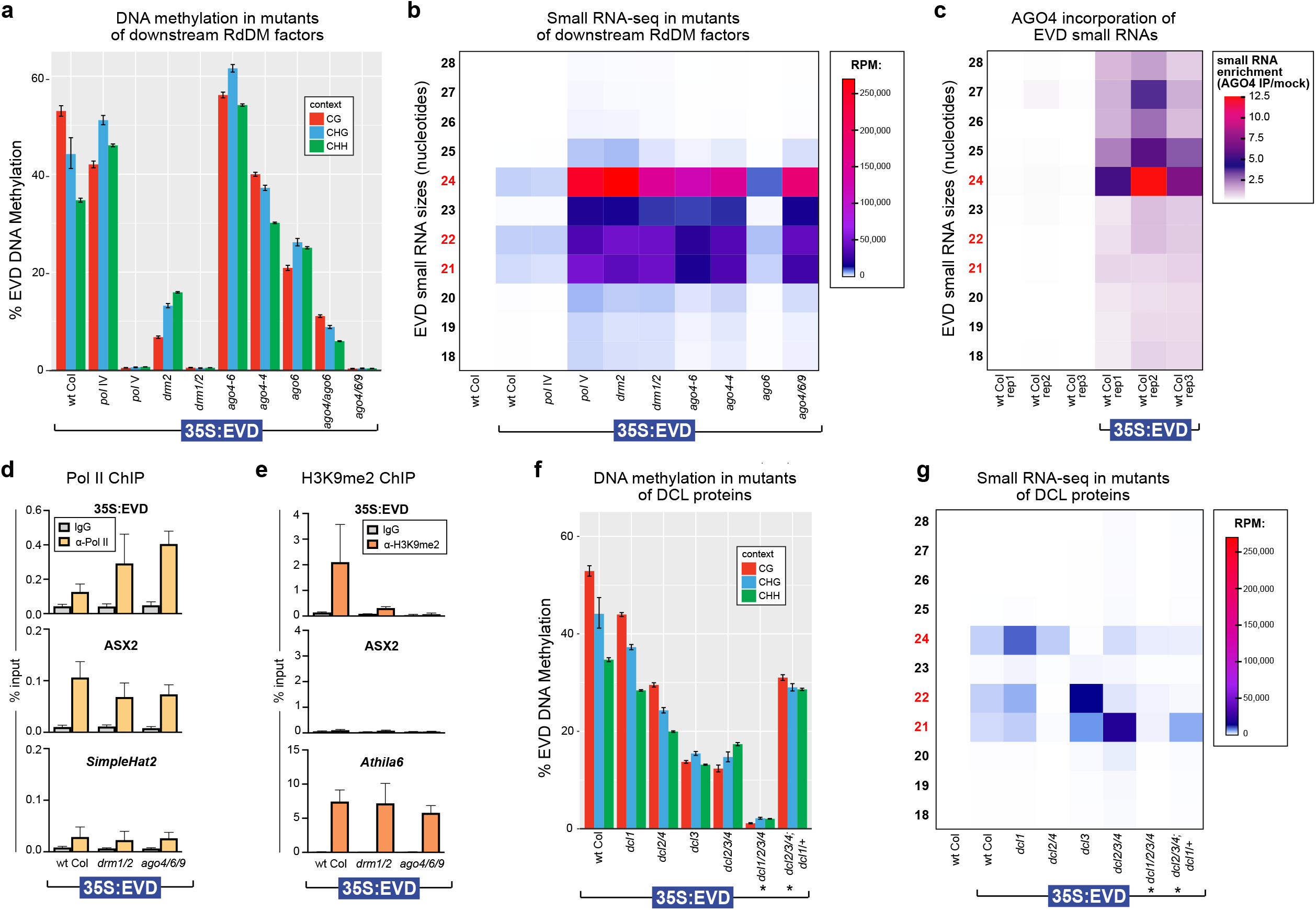
Genetic requirements for the first round of RdDM. A. BSAS of 35S:EVD in T1 plants lacking key RdDM factors demonstrates that the initiation of DNA methylation is dependent upon Pol V, DRM1/2, and at least one member of the AGO4-clade of proteins. B. Heatmap of small RNA sequencing for transgenic lines from part A. The accumulation of different sized siRNAs is shown on the Y-axis, with the key sizes 21, 22 and 24 nt in red. RPM = reads per million genome-mapped sequences. The siRNAs shown were mapped to the entire length of the EVD coding region, not just the region analyzed in part A with BSAS. C. Heatmap of EVD small RNA enrichment in AGO4. Enrichment is calculated as the ratio of small RNA accumulation (RPM) in AGO4-IP over mock-IP samples for each size class of small RNA. D. ChIP analysis of Pol II (Ser5P) in T1 transgenic plants from part A shown as % input. Error bars represent the standard deviation. Three biological replicates were used for each genotype. *SimpleHat2* is a transcriptionally silenced TE negative control, and AXS2 is a transcribed positive control gene. E. ChIP analysis of H3K9me2 levels in T1 transgenic plants shown as % input. Three biological replicates were used for each genotype. *Athila6* is a silenced TE with known accumulation of H3K9me2. F. BSAS of 35S:EVD in T1 plants lacking various combinations of DCL family proteins. The *dcl1/2/3/4* quadruple mutant plants are siblings of the *dcl2/3/4* ; *dcl1*/+ plants (siblings denoted with *). G. Heatmap of small RNA sequencing for the same transgenic lines as in part F.

SiRNA biogenesis is not dependent upon Pol V, DRM1/DRM2 or AGO4/AGO6/AGO9 (Figure 2B, Supplemental Figure 4), confirming that these proteins function downstream of siRNA production during the chromatin-linked phase of RdDM. Instead, mutations in the downstream factors Pol V, DRM1/DRM2 and AGO4-clade proteins surprisingly have increased siRNA accumulation (Figure 2B). We found that this increase in siRNAs positively correlates with the level of Pol II transcription of 35S:EVD in *drm1/2* and *ago4/6/9* mutant (Figure 2D), and is inversely correlated with the level of the transcriptionally-repressive histone mark H3K9me2 (Figure 2E). This demonstrates that in mutants of RdDM downstream machinery (*pol V, drm1/2, ago4/6/9*), without DNA methylation and H3K9me2, 35S:EVD expression is uninhibited, generating more transcription that leads to increased siRNA production. Additional regions of the endogenous genome also generate more siRNAs in *pol V, drm1/2* and *ago4/6/9* mutants (Supplemental Figure 7), suggesting that a lack of downstream RdDM function broadly leads to enhanced Pol II transcription and increased siRNA production.

### Pol II-derived DCL-processed siRNAs trigger RdDM

Our data suggests that siRNA production from 35S:EVD is from Pol II, since siRNA accumulation is not lost in *pol IV* or *pol V* mutants (Figure 2B) and the abundance of siRNAs (Figure 2B) positively correlates with the level of Pol II transcription of 35S:EVD (Figure 2D). In plants, siRNA sizes and categories are determined by the specific DICER family (DCL) protein that produces them^22^. Our data refutes an existing model based on the identical 35S:EVD transgene that RdDM begins only when DCL2 and DCL4 are overwhelmed with double-stranded RNA substrate, thus activating DCL3 to generate 24 nt siRNAs^19^. Instead, we find that *dcl2/4* double mutants result in a ∼50% reduction in DNA methylation (Figure 2F) rather than being hyper-methylated as previously posited^19^. Further, DNA methylation is not entirely dependent upon the presence of 24 nt siRNAs, as *dcl3* mutants that lack 24 nt siRNAs (Figure 2G) do not lose all methylation (Figure 2F)(as in ^16^). Additionally, DNA methylation still persists in *dcl2/3/4* triple mutants, where only DCL1-dependent 21 nt siRNAs remain (Figure 2F-G). These data indicate that 21, 22 and 24 nt siRNAs are all sufficient to trigger the initiation of RdDM. DNA methylation is nearly absent only when siRNAs are severely reduced in *dcl1/2/3/4* quadruple mutants (Figure 2F-G). Therefore, the initiation of DNA methylation is fully dependent on the presence of Pol II-derived DCL-processed siRNAs in a size-independent manner.

### Pol V requires an AGO4-clade protein for localization during the initiation of RdDM

Pol V is essential to establish RdDM (Figure 1), and therefore we aimed to understand how Pol V is recruited to a locus in the absence of preexisting DNA methylation. We re-analyzed published Pol V chromatin immunoprecipitation (ChIP) and RNA immunoprecipitation (RIP) data^12,9^ to determine whether Pol V is present (ChIP) and transcribing (RIP) at low levels throughout the genome. Our re-analysis centered on using mitochondrial genes as additional negative baseline controls, as Pol V subunits are located exclusively in the nucleus^23,24^. We find that Pol V signal is detected at non-RdDM nuclear loci only at the same rate as the mitochondrial negative control genes (Figure 3A). Therefore, rather than patrolling all loci at low levels, Pol V is actively recruited to its new target sites for the initiation of RdDM.

**Figure 3.**
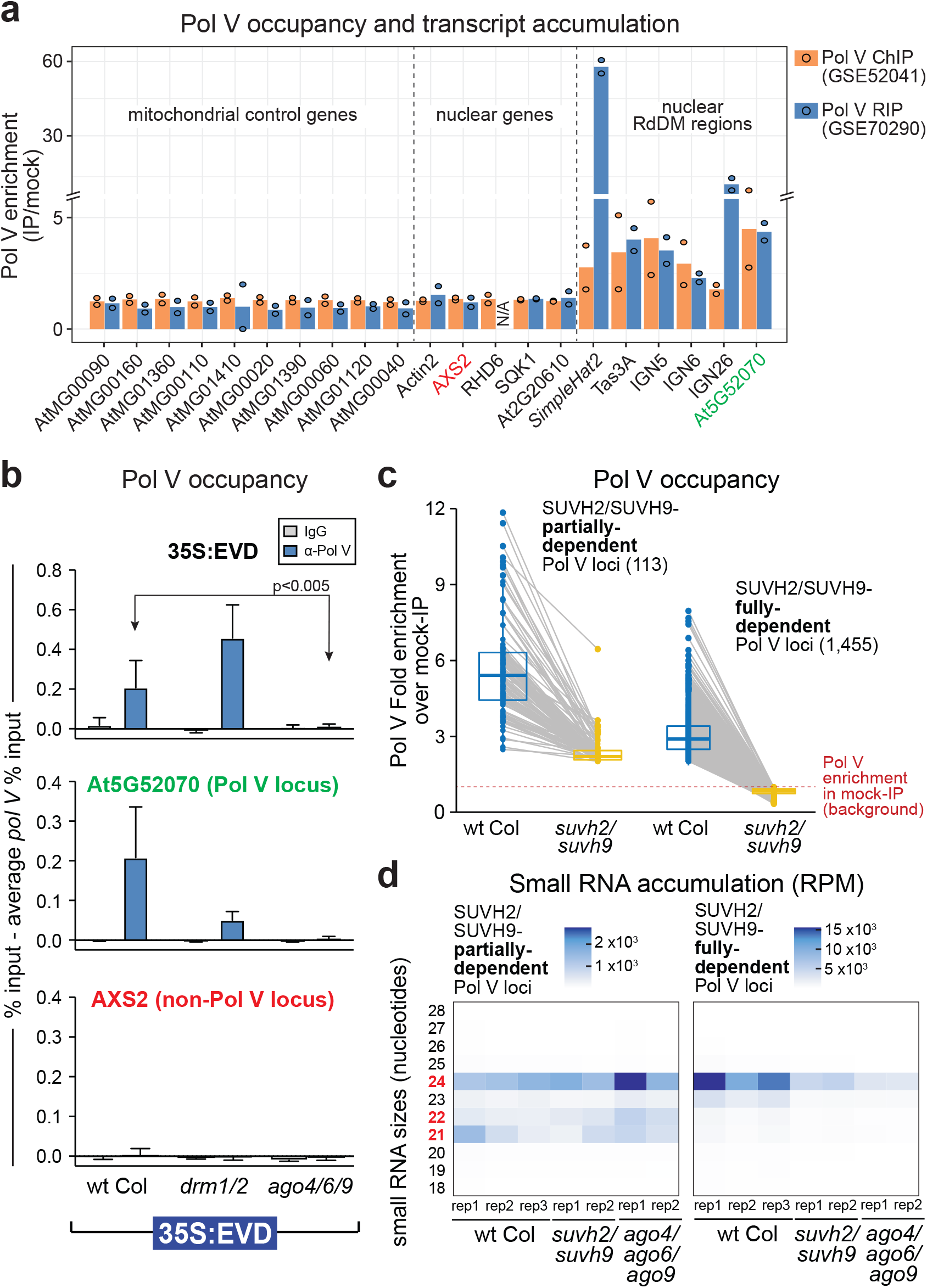
The first round of Pol V recruitment requires an AGO4-clade protein. A. Reanalysis of ChIP and RIP data of Pol V occupancy and transcription. Mitochondrial negative control genes, nuclear genes and nuclear positive control RdDM regions are shown. Two replicates are shown as dots, and their average is the height of the bar. N/A = Not applicable due to absence of any mapping reads in the IP or the mock samples. Genes with colored labels are used as controls in ChIP experiments. B. Pol V ChIP of 35S:EVD T1 plants as % input minus the background of % input in T1 *pol V* mutants. At least three biological replicates were used for each genotype. At5G52070 is a locus that undergoes RdDM and serves as a positive control for Pol V occupancy^32^, while AXS2 is a gene that does not undergo RdDM and serves as a negative control. Error bars represent the standard deviation. Statistical significance was determined by a two-tailed unpaired t-test with Welch’s correction. C. Box plot with connected data points of Pol V occupancy comparing wt Col (blue) and *suvh2/suvh9* double mutants (yellow). Pol V occupied regions are categorized into SUVH2/SUVH9 partially-dependent and fully-dependent loci (see Methods). Pol V occupancy is displayed as fold enrichment over mock-IP. Grey lines represent the change in Pol V enrichment between wt Col and *suvh2/suvh9* for each locus. Box plots represent 25th and 75th percentile with whiskers at 10th and 90th percentile values, and the median is represented by a line. D. Heatmap of small RNA sequencing for Pol V occupied loci that are categorized as either SUVH2/SUVH9 partially-dependent or fully-dependent from part C.

We identified three mutant combinations of downstream RdDM factors that completely lose the ability to target T1 35S:EVD for RdDM (*pol V, drm1/2* and *ago4/6/9*)(Figure 2). To test the requirements of Pol V recruitment to new loci, we performed Pol V (subunit NRPE1) ChIP in these mutant backgrounds in the T1 generation of 35S:EVD plants. We found that the Pol V protein still accumulates in each of these mutant backgrounds (Supplemental Figure 7) and is recruited in the first generation to the TE transgene (Figure 3B). Importantly, this recruitment is not dependent on existing DNA methylation, as Pol V recruitment still occurs in the *drm1/2* mutant (Figure 3B) that lacks DNA methylation (Figure 1). Therefore, this system provides the ability to dissect the methylation-independent recruitment of Pol V. Importantly, we found that Pol V is not recruited for the first round of RdDM in the *ago4/6/9* triple mutant (Figure 3B). This was surprising, as the prevailing dogma suggests that Pol V presence and transcription occurs first and is required to position AGO4-clade proteins at the chromatin target^25^. Conversely, our data demonstrates that AGO4-clade proteins are required to localize Pol V during the initiation of RdDM.

AGO4-clade proteins are directed to their targets by the complementarity of their incorporated siRNAs^26^. To determine if there are regions of the endogenous genome where Pol V is positioned independently of SUVH2/SUVH9 and instead based on AGO4/siRNAs, we first identified regions of the genome where Pol V occupancy is only partially-dependent on the methylation-dependent SUVH2/SUVH9 mechanism (left, Figure 3C). These 113 regions retain siRNA accumulation in *suvh2/suvh9* and *ago4/6/9* mutants (Figure 3D). Conversely, the 1455 regions of the genome where Pol V recruitment is fully dependent on SUVH2/SUVH9 tend to have reduced siRNA accumulation in *suvh2/suvh9* mutants (Figure 3D). These data suggest that when the SUVH2/SUVH9 recruitment method is absent, the continual production of siRNAs is necessary for Pol V occupancy at a small number of loci in the Arabidopsis genome.

### AGO4 can interact with new sites of RdDM independent of Pol V and DNA methylation

We aimed to order the events of recruitment at the chromatin that result in the first round of RdDM. Since Pol V recruitment to 35S:EVD is dependent on a protein of the AGO4-clade (Figure 3), we aimed to determine if the converse was true: Is AGO4’s interaction with chromatin dependent on Pol V? Our data supports this idea that there are a small number of regions of the endogenous Arabidopsis genome where AGO4 interacts with target chromatin loci independent of a Pol V-derived scaffolding transcript. Since AGO4 targeting is dependent on siRNA complementarity, we began by identifying loci that continue to produce 23-24 nt siRNAs in a *pol V* mutant (Figure 4A). We overlapped these 4246 Pol V-independent siRNA loci with 820 previously-identified AGO4 bound loci^27^, resulting in 91 testable AGO4-bound regions of the genome that do not lose siRNAs (Figure 4B). 63 of these testable AGO4-bound regions (69%) retain AGO4 occupancy in the *pol V* mutant (Figure 4C), demonstrating AGO4 recruitment without Pol V.

**Figure 4.**
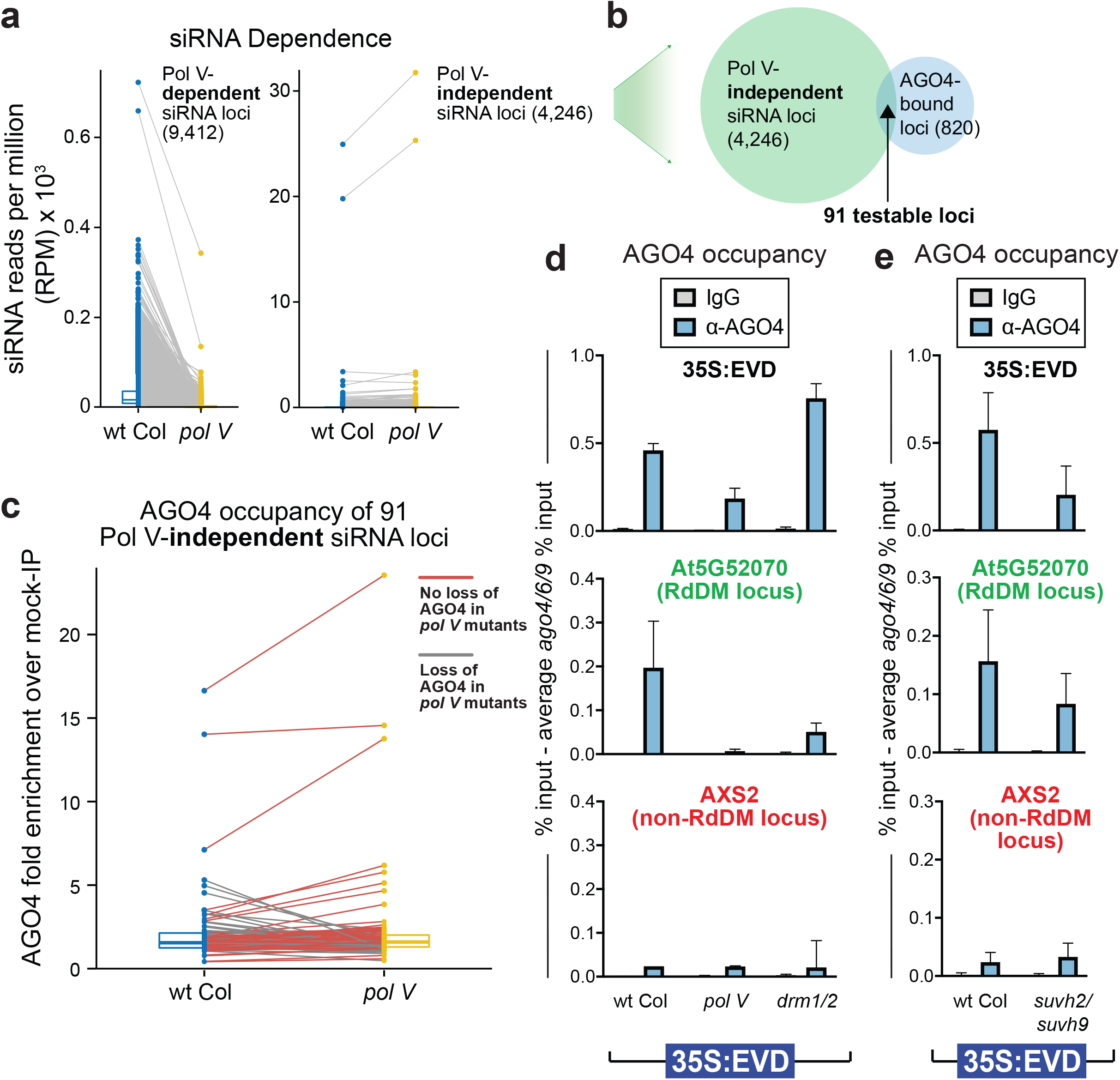
AGO4 can localize to chromatin loci independent of Pol V. A. Box plot with connected data points of 23-24 nt siRNA accumulation in wt Col (blue) and *pol V* mutants (yellow). SiRNA loci are defined as Pol V-dependent or -independent based on change in siRNA accumulation in *pol V* compared to wt Col. Box plot percentiles are the same as in Figure 3C. B. Venn diagram showing the overlap of Pol V-independent siRNA loci from part A and AGO4-enriched loci as previously defined^27^. The overlap provides 91 testable loci for part C. C. Box plot with connected data points of AGO4 protein enrichment. AGO4 occupancy is displayed as fold enrichment over mock-IP. Grey lines display loci that lose AGO4 occupancy in *pol V* mutants (31%). Red lines display loci that retain AGO4 occupancy in the *pol V* mutant background (69%). D. AGO4 ChIP of 35S:EVD T1 plants as % input minus the background of % input in T1 *ago4/6/9* mutants. Bar graph display is the same as in Figure 3B. At5G52070 is a positive control locus that undergoes RdDM, and AXS2 is a negative control gene that does not go through RdDM. E. AGO4 ChIP of 35S:EVD T1 plants as % input minus the background of % input in T1 *ago4/6/9* mutant plants. Bar graph display is the same as in Figure 4D.

To determine if AGO4 is localized to the 35S:EVD transgene during the initiation of RdDM, we performed AGO4 ChIP on T1 35S:EVD plants. As a control, we confirmed that the AGO4 protein is present in the various mutants we tested (Supplemental Figure 7). In the ChIP experiment, we detected AGO4 at 35S:EVD during the initiation of RdDM in wild-type plants, as well as in *pol V, suvh2/suvh9* and *drm1/2* mutants (Figure 4D-E). The level of AGO4 at 35S:EVD in *pol V* and *suvh2/suvh9* mutants is not as high as in wt Col plants, but is nonetheless significantly higher than the negative control AXS2 gene and the IgG negative control (Figure 4D-E). In addition, the AGO4 ChIP signal in the *drm1/2* mutant may be higher due to the increased siRNAs produced in this mutant (Figure 2B). We conclude that AGO4 is directed to the target transgene without the requirement of a Pol V scaffolding transcript or DNA methylation, and is independent of the known SUVH2/SUVH9 recruitment mechanism of Pol V (Figure 4E), placing AGO4 interaction with the target locus before and not reliant upon Pol V activity.

### siRNAs are sufficient to direct Pol V function

Since AGO4 can be recruited to a new target locus independent of Pol V, we aimed to determine if siRNAs are sufficient to direct Pol V activity and the first round of RdDM. Our 35S:EVD transgene system cannot address this question, since the source locus of the siRNAs and the target of Pol V action are the same (*cis*-acting). Instead, we generated a *trans*-acting two component system to uncouple siRNA production from Pol V recruitment. SiRNAs are generated from an inverted repeat transgene that targets an unmethylated endogenous gene in *trans* (Figure 5A)(as in ^28,29,30^). The endogenous gene we targeted (At3G12210) encodes a broadly expressed DNA-binding protein that does not produce siRNAs (Figure 5B), is not methylated (Figure 5C), and is not bound by Pol V (Figure 3A) in wt Col plants. We named this uncharacterized gene SQUEAKY1 (SQK1). Upon addition of the SQK1 inverted repeat transgene (‘SQK1-IR’), 21, 22 and 24 nt siRNAs accumulate from the hairpin region (Figure 5B), and these siRNAs function in *trans* to direct RdDM to the SQK1 endogenous gene (Figure 5C-D). In a *pol V* mutant, siRNAs are still generated from the IR transgene (Figure 5B), but RdDM does not occur (Figure 5C-D), again demonstrating that Pol V acts downstream of siRNA production and is essential for RdDM. As with 35S:EVD, the initiation of RdDM is fully dependent on an AGO4-clade protein (Figure 5C-D). Taken together, these data demonstrate that the production of new siRNAs and the presence of an AGO4-clade protein are sufficient to target Pol V-dependent RdDM to a new non-transgenic locus independent of existing methylation.

**Figure 5.**
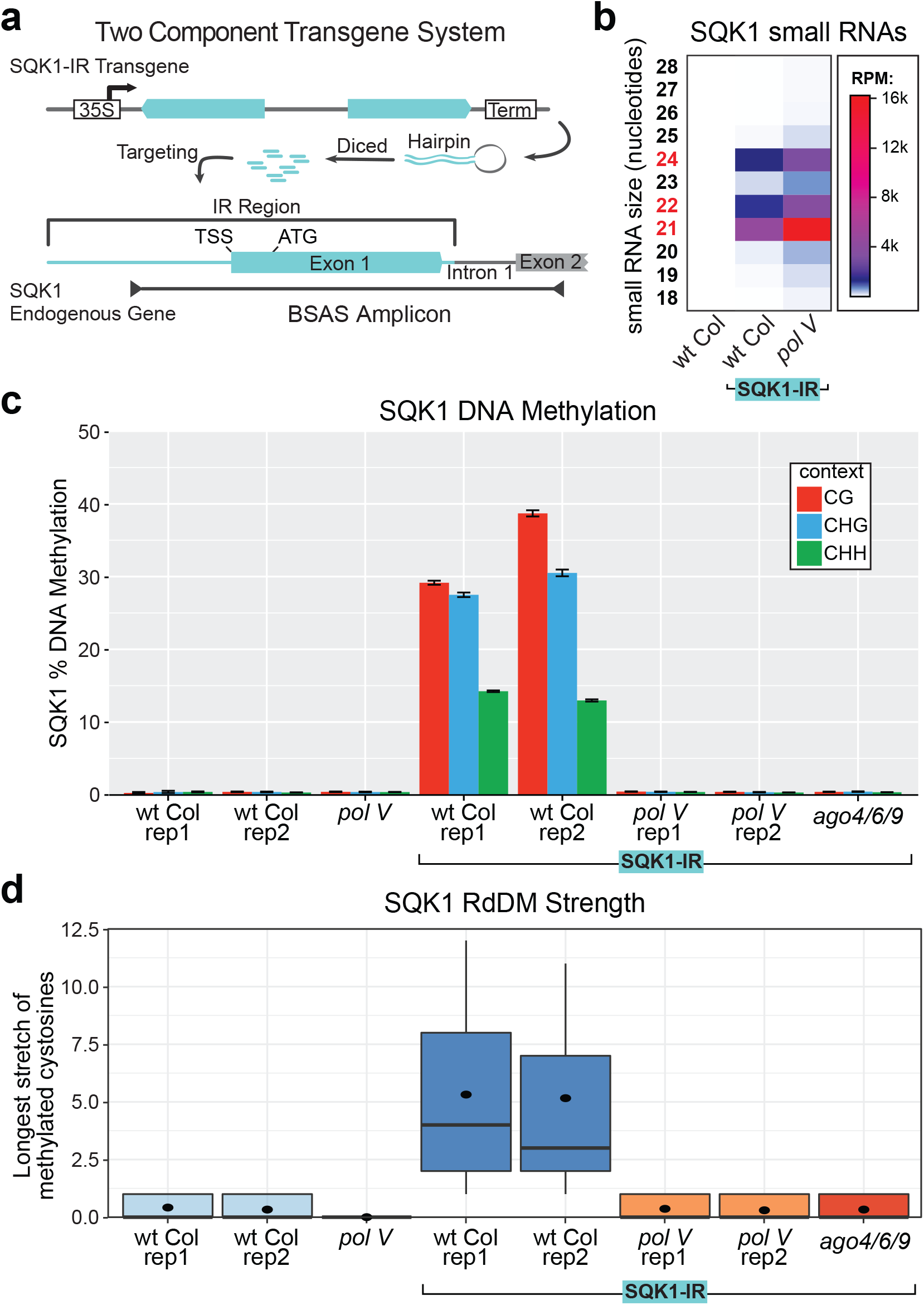
siRNAs are sufficient to target the first round of RdDM. A. Two component system with siRNA production from a T1 inverted repeat (IR) transgene targeting the endogenous gene SQK1 for RdDM. B. Heatmap of small RNA sequencing. C. BSAS for T1 transgenic SQK1-IR plants. D. Box plots of RdDM strength for plants from C.

### Pol II expression is necessary for permitting the first round of RdDM

While Pol II cannot substitute for Pol V (Figure 1), we have identified that Pol II does play a role in making the target locus receptive to the first round of RdDM. While investigating the targeted methylation of the SQK1 gene (Figure 5), we observed that the CHH methylation (indicative of RdDM) was not distributed evenly across the region targeted by siRNAs. Instead, methylation peaks upstream of the transcriptional start site (TSS) near the proximal SQK1 promoter (Figure 6A). This pattern of methylation does not correlate with the abundance of siRNAs across this region (Figure 6A), suggesting that the observed methylation pattern is based on the SQK1 promoter activity. To test the role of Pol II expression during the first round of RdDM, we used Cas9 to generate a 1298 bp deletion of the SQK1 promoter (*sqk1-1*)(shown in Figure 6A). This severely reduces, but does not completely eliminate SQK1 expression (Figure 6B). Where on the *sqk1-1* allele Pol II initiates and the direction of transcription is not known. When the homozygous *sqk1-1* mutation is combined with the SQK1-IR, although the siRNAs still target *sqk1-1*, methylation is significantly reduced (Figure 6A,C). In a separate test, we generated a second IR targeting system (as in Figure 5A) for a different gene that has a specific and limited developmental expression pattern (Supplemental Figure 8). Our data for both two-component IR systems (Figure 6A-C and Supplemental Figure 8) demonstrates that the strength of RdDM positively correlates with the level of Pol II expression at the target locus.

**Figure 6.**
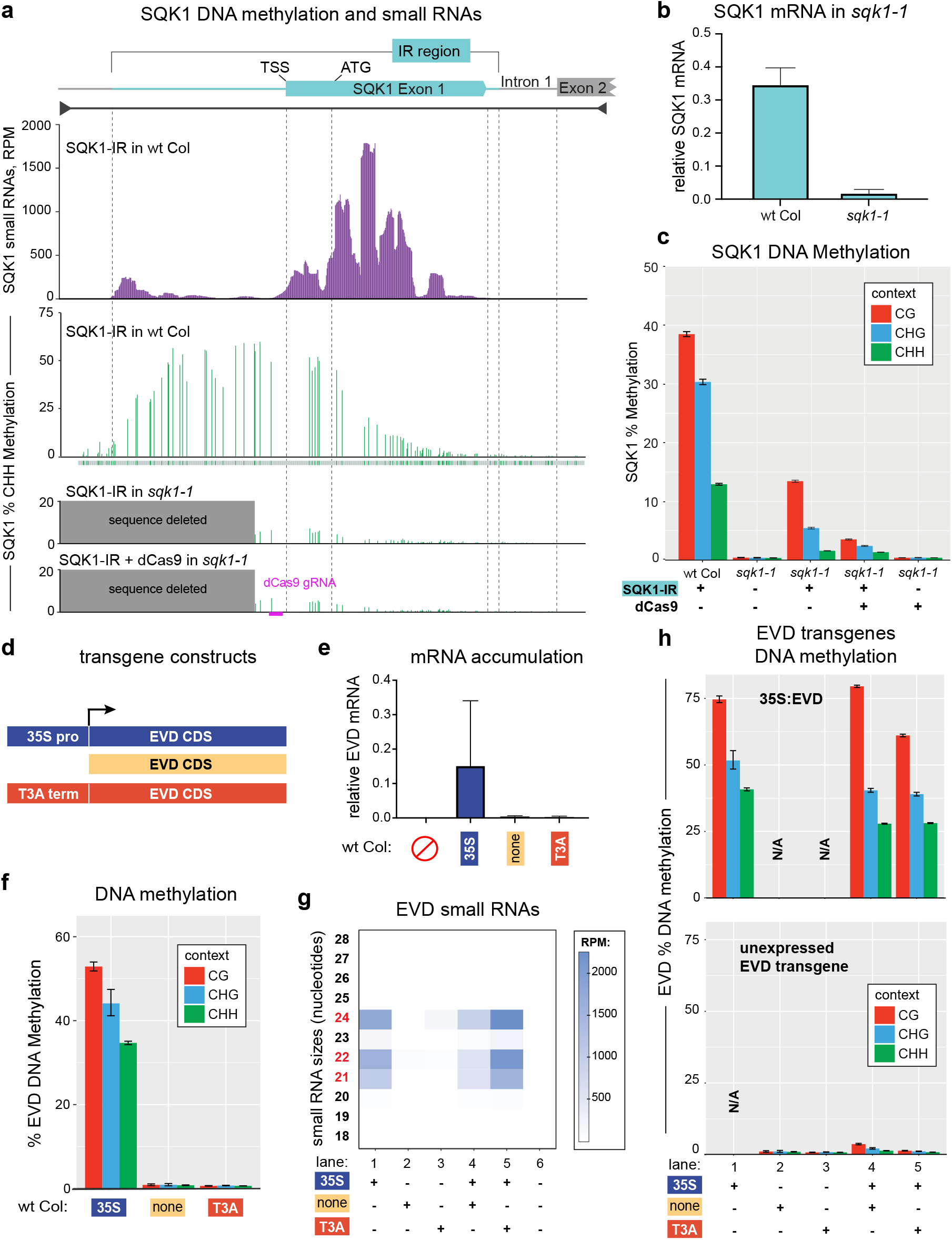
Pol II’s role in the first round of RdDM. A. CHH context DNA methylation and siRNAs (combined 21, 22 and 24 nt) aligned to the endogenous SQK1 gene. The region of SQK1 that matches the SQK1-IR transgene is annotated. The track below the SQK1-IR of wt Col CHH methylation identifies where each CHH context cytosine is located (green), even if unmethylated. The DNA methylation track for SQK1-IR in wt Col is produced from BSAS of two overlapping amplicons. Dashed lines align key annotation features of the SQK1 gene, and the location of the dCas9 gRNA sequence is annotated. B. qRT-PCR of plants homozygous for the SQK1 promoter deletion allele *sqk1-1*. The error bar represents the standard deviation of three biological replicates. C. BSAS of plants with and without the SQK1-IR, the homozygous *sqk1-1* deletion and dCas9. D. Depiction of three EVD TE transgenes. These transgenes share the exact coding sequence of EVD5 (AT5TE20395) lacking any vector-encoded terminator sequence while differing in their upstream elements. These upstream elements include the 35S constitutive promoter (from Figure 1), no promoter and the T3A terminator. E. qRT-PCR of EVD mRNA in T1 transgenic plants indicates that only 35S:EVD has Pol II expression. F. BSAS of T1 transformants for each respective EVD transgene in wt Col. G. Heatmap of small RNA sequencing from single and double EVD transgenic plants. The siRNAs shown were mapped to the entire length of the EVD coding region, not just the region analyzed by BSAS in part F. H. BSAS for the 35S:EVD transgene (top) and the second unexpressed EVD transgene (bottom) in the same transgenic individuals. N/A = not applicable, as that transgene is not in this plant line. Biological replicates are shown in Supplemental Figure 8.

To decisively test the necessity of Pol II expression in the first round of RdDM, we created two new EVD transgenes that are definitively either expressed or not. These include the ‘EVD-only’ (EVD coding sequence with no promoter) and ‘T3A-EVD’ transgenes (T3A terminator directly 5’ of the EVD coding sequence, to ensure no read-through Pol II transcription)(Figure 6D). qRT-PCR of EVD transgenes confirms that only 35S:EVD has appreciable mRNA accumulation (Figure 6E), although the expression levels generated by the 35S promoter are highly variable due to the nature of T1 transgenesis^31^. We confirmed that RdDM of 35S:EVD is dependent on Pol II expression, as only the expressed transgene version is targeted for methylation (Figure 6F, biological replicates in Supplemental Figure 8). The key test was when we combined expressed and unexpressed EVD transgenes in the same genome. We found that whenever the expressed 35S:EVD transgene is present, it produces abundant siRNAs (Figure 2B and Figure 6G lanes 1, 4, 5), which are incorporated into AGO4 (Supplemental Figure 6) and drive RdDM to 35S:EVD itself (Figure 1 and Figure 6H lanes 1, 4, 5). When a second promoterless EVD transgene is introduced into the same plant genome that has 35S:EVD, these unexpressed transgenes do not become methylated (Figure 6H lanes 4-5, biological replicates in Supplemental Figure 8). These double-transgenic plants have EVD siRNAs that are incorporated into AGO4 and are competent to perform RdDM (evidenced by the fact that 35S:EVD is targeted by RdDM in the same plant (Figure 6G lanes 4-5)). The only difference as to why one transgene is methylated and the other is not is the Pol II activity at the target transgene. Therefore, Pol II transcription is not sufficient to substitute for the absence of Pol V, but Pol ll activity is necessary to make a locus receptive to the first round of RdDM.

### Pol II’s role in the initiation of RdDM

We aimed to identify the exact function of Pol II during the first round of RdDM. Pol II could be involved in producing a scaffolding transcript for AGO/siRNA interaction. Alternatively, AGO/siRNA complexes are known to interact with single-stranded DNA^32^. Pol II’s function at the target locus could be to open the double-stranded structure of the DNA, allowing for the AGO/siRNA complex to base pair with single-stranded DNA, which fits the theory that RdDM is focused at DNA replication forks^33^. To separate these models, we attempted to increase the RdDM of the low-expressed *sqk1-1* allele by targeting dCas9 to this locus. We used a gRNA to target dCas9 and R-loop formation^34^ to the homozygous *sqk1-1* locus in the presence of the SQK1-IR (see gRNA location in Figure 6A). This did not result in higher methylation (Figure 6A,C) as would be expected if DNA opening was the RdDM function of Pol II at the target locus. Instead, methylation decreased, as though dCas9 interferes with the low level of Pol II transcription at this locus. Since this dCas9 experiment provided negative results, several alternative interpretations exist. However, this experiment suggests that the function of Pol II at a new RdDM target locus is likely to generate the RNA transcript that acts in AGO/siRNA complementary base pairing.

## Discussion

The roles of RNA Polymerase IV and V at silenced TE fragments undergoing the self-reinforcing cycle of RdDM are well described^35^. However, the specific roles of these polymerases and Pol II during the initiation phase of TE and transgene silencing have remained enigmatic. Here we dissect polymerase function using newly integrated transgenes. This strategy permits the discrimination of the first round of RdDM from those already engaged in the RdDM cycle.

We find that Pol V is required for all *de novo* RdDM, and Pol II cannot substitute for this function. Our data refutes the model whereby Pol II transcripts are able to initiate the first round of DNA methylation. The essential function of Pol V during the initiation of RdDM correlates with the recent finding that Pol V is a key factor in the evolutionary repression of TEs in the Arabidopsis lineage^36^. Our data also refutes the ‘Pol V surveillance’ model, whereby Pol V produces scaffolding transcripts that, when an siRNA is present, will trigger the first round of methylation. It has been recently demonstrated that the artificial tethering of a Pol V-recruitment factor to a genomic locus triggers RdDM^37^. Therefore, if Pol V surveilled everywhere, we would expect to observe spurious low-level methylation across the genome, which is not detected, even in DNA glycosylase mutants that fail to remove DNA methylation from the genome^38^. Although our data refutes the Pol V surveillance model, our findings agree with the point that siRNAs are the key determinant to target new RdDM^15^.

Our data demonstrates a methylation-independent mechanism for recruitment of Pol V to a new target locus. Previous reports suggested that Pol V is recruited independently of AGO4 and siRNAs, and RdDM would occur only when both Pol V and AGO4 were independently recruited to a locus^39^. Our results suggest a more linear ordered pathway of recruitment, beginning with AGO4 directed by siRNAs. We find that AGO4 is recruited to a new site of RdDM independent of and before Pol V and DNA methylation, and AGO4-clade proteins are themselves guided by the complementarity of the incorporated siRNA. We propose an siRNA-directed pathway of Pol V recruitment in Figure 7.

**Figure 7.**
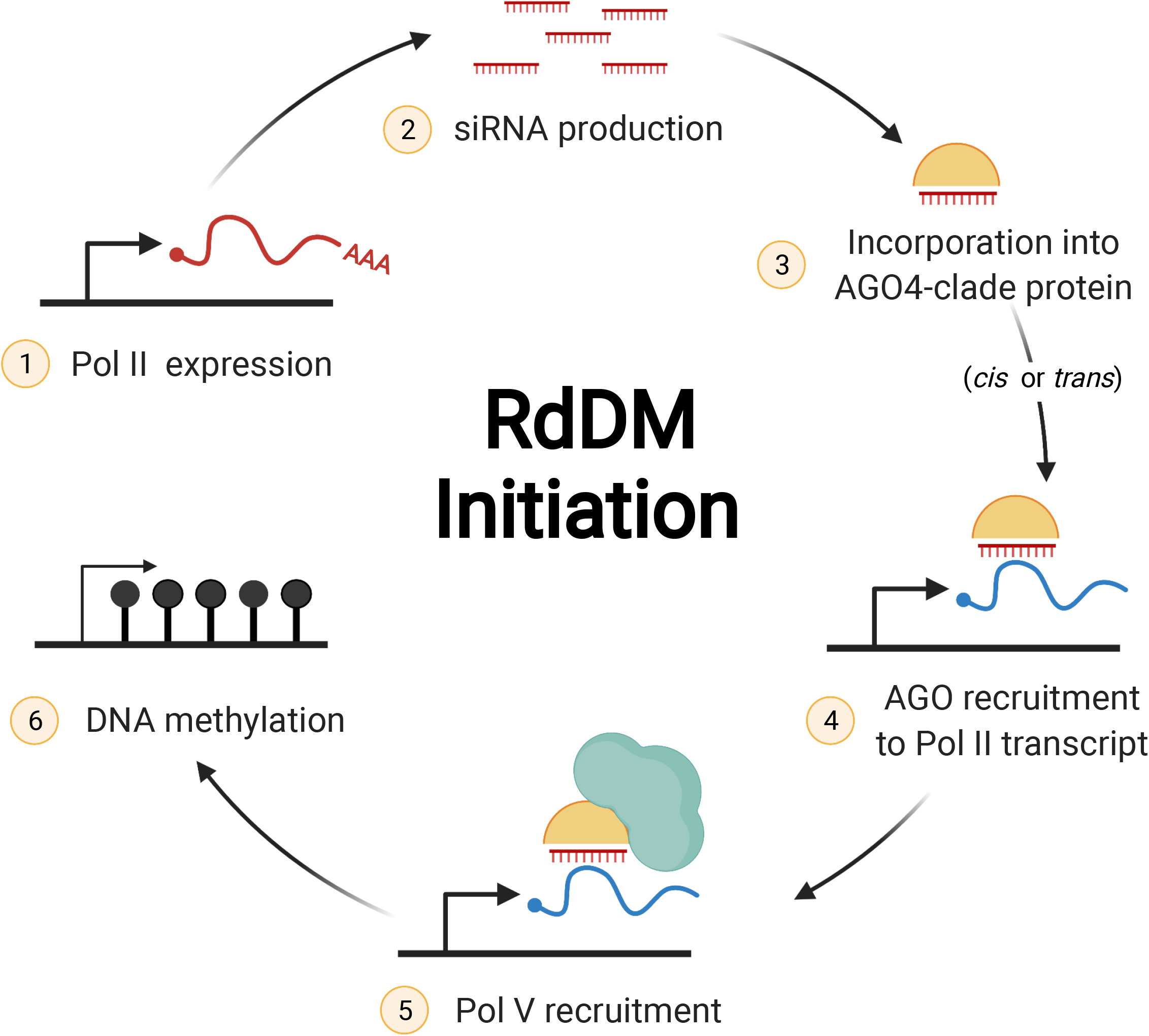
Model of RNA-driven recruitment of Pol V to new RdDM targets. The initiation of RdDM begins with Pol II expression of the unmethylated locus (step 1), followed by siRNA production from that transcript by any member of the DCL family (step 2). SiRNAs are incorporated into a member of the AGO4-clade of proteins (step 3) and this can only target RdDM to complementary regions of the genome that are expressed (step 4). *Cis* action refers to a region that generates siRNAs and targets its own methylation, while in *trans* action the siRNAs are acting on a separate locus. The AGO4 targeting of a Pol II expressed region of the genome results in Pol V recruitment (step 5) and the first round of RdDM (step 6). This final methylated product can then enter the methylation-dependent pathway of continual Pol V recruitment guided by SUVH2/SUVH9. See Discussion section for details. Created with BioRender.com.

Pol II’s function during the first round of RdDM is two-fold. First, it is necessary to produce the raw transcripts for siRNA production (step 1, Figure 7). During the self-reinforcing cycle of RdDM at endogenous silenced TEs, this function is taken over by Pol IV. However without existing heterochromatic marks to recruit Pol IV, the new region must be transcribed by Pol II to generate siRNAs in *cis* or siRNAs provided from a separate locus in *trans*. We find that in cases where Pol II is generating siRNAs, mutations in the downstream RdDM factors (such as *pol V, drm1/2* and *ago4/6/9*) result in the lack of DNA and H3K9 methylation, leading to more Pol II expression and more substrate for the higher accumulation of siRNAs (Figure 2). Once the raw transcripts are produced by Pol II, we find that they can be processed by any DCL protein into 21-24 nt siRNAs (step 2, Figure 7). All of these siRNAs sizes, generated by any of the four Arabidopsis DCL proteins, are capable and sufficient to target the first round of RdDM. Our data agrees with the ‘saturation’ model whereby during the initiation of RdDM all available DCL proteins function to process the large volume of Pol II transcripts, producing 21, 22 and 24 nt siRNAs that are all capable of guiding RdDM^16^. These siRNAs are loaded into an AGO4-clade protein (step 3, Figure 7), as RdDM function was abolished in the *ago4/6/9* triple mutant.

Second, Pol II action is required downstream of siRNA production. Even when complementary siRNAs and AGO4 are present, the first round of RdDM is dependent upon Pol II activity at both transgenes and endogenous genes (Figure 6). Our data suggests that the role of Pol II at the target locus is to generate the first scaffolding transcript for AGO/siRNA interaction (step 4, Figure 7). This is supported by the protein-protein interaction of the Pol V recruitment factor RDM1 with Pol II, AGO4 and DRM2^24^. This function of Pol II suggests that it retains some of its ancestral ability to generate scaffolding transcripts, which has otherwise been subfunctionalized and relegated to Pol V. Even though the Pol II transcript interaction with the AGO/siRNA complex does not result in methylation, it is necessary for the recruitment of Pol V (step 5, Figure 7) and subsequent RdDM (step 6, Figure 7). A recent study demonstrated that Pol II’s C-terminal domain can act to recruit more Pol II protein^40^, and during the initiation of RdDM Pol II may act in a similar manner to recruit Pol V, target RdDM and result in the establishment of epigenetic inheritance.

## Methods

### Plant growth and lines

*Arabidopsis thaliana* plants were grown at 22°C on Pro-Mix FPX soil in Conviron MTPS-120 growth chambers with 16-hour 200 µmol/m^2^/s light. The specific alleles of all mutants are shown in Supplemental Table 1. Inflorescence (flower buds, stage 1-12) tissue was used for all experiments unless otherwise noted. All transgenic material is the first generation (T1) after integration of the T-DNA into the genome unless otherwise stated. All transgenic lines were stably integrated and produced by the Agrobacterium-mediated floral dip method and subsequent selection for Basta- or Hygromycin-resistant plants. Biological replicates are non-overlapping pools of tissue collected from T1 transgenic plants. Double EVD transgenic lines (Figure 6G-H) were generated by floral dipping T2 Basta-resistant 35S:EVD plants with Hygromycin-resistant EVD-only and T3A-EVD transgenes. Screening and genotyping in the next generation resulted in T3 35S:EVD + T1 EVD-only or T1 T3A-EVD plants. To create the *dcl1/2/3/4* 35S:EVD transgenic plants, we transformed a line that was homozygous *dcl2/3/4* and heterozygous for the *dcl1-9* allele. Resulting transformants were genotyped for DCL1, and divided into those that were homozygous mutants and those that were DCL1 heterozygotes.

### Transgene Generation

Transgenes were generated from primers listed in Supplemental Table 1. The EVD coding sequence (At5TE20395) was directly amplified from wt Col DNA using primers containing appropriate plasmid homology for In-Fusion Cloning (Clontech) into their respective digested binary vector backbones. Both pB2GW7 (Basta selection) and pH2GW7 (Hygromycin selection) digested with SpeI/HindIII were used for 35S:EVD, as some mutants that were transformed with this transgene already contain a Basta resistance cassette. pH2GW7 digested with SacI/HindIII was used for EVD-only. For T3A-EVD, the T3A terminator and EVD coding sequence were separately amplified and then joined together using overlapping PCR before In Fusion Cloning into pH2GW7 digested with SacI/HindIII.

The inverted repeat regions of transgenes were synthesized by Thermo-Fisher and cloned into the binary vector pEG100 via SpeI/XhoI restriction digest followed by ligation using T4 DNA Ligase (NEB). The SQK1 inverted repeat contains bases AtChr3 3895083:3894528 (order reversed to prevent open reading frame) followed by the PDK intron and then AtChr3 33894528:3895083 (resulting plasmid named pMJS064). The RHD6 inverted repeat was fashioned similarly containing bases AtChr1 24795653:24795105. See Supplemental Table 1 for inverted repeat sequences. A Hygromycin-resistant version of each of these transgenes was created by amplifying the 2X 35S promoter: Hygromycin b phosphotransferase: 35S terminator cassette from the pDIRECT_21B vector using primers from Supplemental Table 1 to add MluI and SacI restriction sites before being digested and ligated into the IR-pEG100 vectors in place of the Basta cassette.

The *sqk1-1* allele was created using the egg-cell promoter/enhancer CRISPR-Cas9 system described by ^41^. The two gRNA cassette was created using a four primer overlapping PCR (see Supplemental Table 1) to amplify the 20 nt gRNA : U6-26 terminator : U6-29 promoter : 20nt gRNA from the pCBC-DT1T2 plasmid^42^. This PCR product was cloned via BsaI Golden Gate reaction into a version of the pHEE401E plasmid containing an additional NapA promoter : dsRED : Nos terminator at the MfeI restriction site. The dsRED cassette was amplified out of the Traffic Lines^43^ (primers in Supplemental Table 1) and provides a method for negative selection of the Cas9 transgene. T1 transformants were selected on Hygromycin plates and screened for SQK1 promoter deletion via PCR (primers in Supplemental Table 1).

To create the dCas9 + SQK1-IR system, the pHEE401E plasmid was digested with NcoI and EcoRI to remove the active Cas9 cassette. The dCas9 sequence was amplified from the pDIRECT_21B plasmid^44^. The Rps5a promoter for dCas9 was synthesized by ThermoFisher and cloned into the pHEE401E backbone via In-Fusion reaction (Clontech). A gRNA cassette was created and inserted via Golden Gate reaction as described above using primers in Supplemental Table 1. dCas9 targets SQK1 at the sequence ATACTCAAAAATTAATAGA. The gRNA - dCas9 complex was amplified (primers in Supplemental Table 1) and In-Fusion cloned (Clontech) into a SacI digested pMJS064 (described above) that contains the SQK1-IR. The Rbs terminator for dCas9 was amplified from pHEE401E and In-Fusion cloned into the pMJS064 + dCas9 plasmid following digestion with AvrII (resulting plasmid named pMJS082). The dCas9 construct without the SQK1-IR was created by removing the SQK1-IR from pMJS082 via restriction digest with SpeI and XhoI.

### Expression Analysis

RNA was extracted from three biological replicates using TRIzol Reagent (Thermo Fisher) and then DNase I treated and cleaned-up with the TURBO DNase DNA-free kit (Thermo Fisher) according to manufacturer’s protocol. cDNA synthesis was performed using an oligo d(T) primer and Tetro Reverse Transcriptase (Bioline). We performed qRT-PCR using the primers in Supplemental Table 1 and SYBR green supermix. Relative expression was determined using the 2 ^-ΔΔCt^ method, comparing the gene of interest to a housekeeping control gene (AXS2 or ACTIN 2, see Supplemental Table 1). Mean and standard deviations of 3 biological replicates are calculated by Graphpad Prism and shown as bar graphs with error bars.

### Bisulfite Amplicon Sequencing and Analysis

Genomic DNA was treated with RNase A, then cleaned and recovered by phenol chloroform isolation. 500 pg - 2 µg DNA was treated with sodium bisulfite using the EZ DNA Methylation kit (Zymo Research). Amplicons for sequencing were generated by PCR using degenerate primers (see Supplemental Table 1) for each locus using My Taq HS mix (Bioline) and gel purification. For each bisulfite conversion reaction performed, we also performed PCR on a control unmethylated locus (At2G20610) in order to calculate the conversion rate (see Supplemental Figure 1 for example).

To generate amplicon libraries, purified amplicons were pooled in equimolar ratios. 100 ng of purified amplicon DNA in 30 µL volume were used for library preparation with the Illumina Nextera DNA Flex kit. BSAS libraries from December 2019 to present were created using a modified Nextera Flex protocol (first described by ^45^). Per index, 1 µL Tagmentation (BLT) beads from the Nextera Flex kit was diluted with 19 µL of ultrapure water and combined with 30 µL sample DNA and 50 µL lab-made 2X tagmentation buffer (20 mM Tris, 20 mM MgCl, 50% DMF) before being tagmented at 55°C for 15 minutes. Tagmenation was stopped by adding 20 µL 0.2% SDS and incubating at 37°C for 15 minutes. Beads were washed three times with 100 µL lab-made tagmentation wash buffer (10% PEG8000 and 0.25 M NaCl in TE buffer).

Tagmented DNA was amplified directly from the beads via 6 cycles of PCR amplification using the PrimeSTAR GXL Polymerase kit (Takara) and dual indexed adapters (Illumina) in 45 µL reactions. Finally, libraries were purified using 81 µL SPRIselect beads (Beckman Coulter) or Nextera Purification Beads (Illumina), washed twice in 200 µL 80% EtOH and eluted off beads in 32 µL ultrapure water.

We sequenced the resulting libraries with 300 nt single-end reads on the Illumina MiSeq platform at the University of Delaware DNA Sequencing and Genotyping Center. Raw reads were trimmed for adapters and mapped to all the amplicons pooled together for sequencing using *methylpy* (https://github.com/yupenghe/methylpy)^46^. Reference DNA sequences have primer sequences removed. The allC files (see Data and Code Availability) containing methylation data for each cytosine were used as input for Bedtools ^47^ to calculate methylation percentage for each locus and plotted using ggplot2 in R. 95% Wilson confidence intervals were calculated as in ^48^ for error bars. BSAS data of DNA methylation levels was quantified per amplicon as the average methylation at each cytosine sequence context (CG, CHG, CHH). The total number of reads reporting the methylation status of a locus is noted as BSAS coverage. Data was analyzed in R and plotted with ggplot2.

### Bisulfite Sanger Sequencing and Analysis

Amplicons for Sanger sequencing from Supplemental Figure 1 were generated as above for BSAS. Purified amplicons were subjected to single colony purification by TOPO TA cloning into pCR4 (Thermo Fisher) and transformation into *E. coli*. Individual colonies were sequenced by Sanger sequencing (Eton Biosciences) and analyzed in *Kismeth*^*49*^ using default parameters.

### Analysis of Whole Genome Bisulfite Sequencing Data

Genome-wide MethylC-seq data is publicly available for wt Col Arabidopsis inflorescence^50^. Processed data was downloaded from GEO (GSM2101949). Similar to BSAS analysis, the allC file was used as input for *Bedtools*^*47*^ to calculate methylation percentage for each locus shown in Supplemental Figure 1.

### Determination of RdDM Strength

To measure RdDM strength, we calculated the number of consecutive cytosines that are methylated for a given locus per sequencing read. Using *methylpy*^*46*^ we isolated the reads that mapped to a specific locus and used these as input for *Kismeth*^*49*^. The *Kismeth* output displays an image of methylation status of each individual cytosine along each individual read, and this was used for image analysis. We used a custom python script (https://github.com/jpeasari/Dot-Plot-Anaysis-OpenCV) to analyze each read in the image, represented by one row of methylation data and determined the longest stretch of consecutive methylated cytosines in each row. We summarized the longest stretch counts for each locus in box plots using ggplot2 in R.

### Small RNA Sequencing and Analysis

Small RNA was sequenced as in ^17^. Briefly, Trizol reagent (Thermo Fisher) was used to isolate total RNA. The mirVana miRNA isolation kit (Thermo Fisher) was used to enrich small RNAs. The TruSeq Small RNA Library Preparation kit (Illumina) was used to generate libraries and multiplexed for sequencing on a HiSeq 4000, HiSeq X or NextSeq 550 system at the University of Delaware DNA Sequencing and Genotyping Center or Novogene Inc.

Post-sequencing, the Illumina universal adapter was removed from the demultiplexed libraries using the fastx toolkit. Total genome matching reads were calculated using *bowtie* (parameters: -v 0), and this value is used for library depth normalization. The *sRNA Workbench*^*51*^ was used to filter out t/rRNA reads, low complexity reads and retain only 18-28 nt small RNA reads that map the Arabidopsis Araport11 genome. To map the small RNAs to the genome, *ShortStack*^*52*^ was used with parameters: --nohp --mmap f --bowtie_m all --align_only. For assaying siRNAs from a specific transgene, *bowtie* (parameter: -v 0 --best --strata -M 1) was used to map the small RNAs to the full transgene sequence. *Bedtools* was used to count the number of reads mapping a specific locus. In Figure 3D, replicate 2 of *ago4/6/9* contains the 35S:EVD transgene (from Figure 2B). The small RNAs mapping the transgene were removed before analysis for Figure 3D. In Figure 4A, *de novo* clusters of 23-24 nt siRNAs were called using *ShortStack* with a threshold of at least 10 raw reads in wt Col and Pol V-dependence was determined by at least a ≥2-fold loss of siRNAs in *pol V* mutants. Conversely, loci that did not lose siRNAs in *pol V* compared to wt Col were categorized as Pol V-independent siRNA loci. ggplot2 in R was used to generate siRNA heatmaps.

### AGO4-incorporated small RNA library preparation and analysis

Frozen inflorescence tissue was ground with liquid nitrogen and resuspended in lysis buffer (50 mM Tris pH 8, 150 mM NaCl, 5 mM MgCl_2_, 10% glycerol, 1% IGEPAL, 0.5 mM DTT, 1mM PMSF, 1X Roche protease inhibitor cocktail) and homogenized with mixing for 15 minutes at 4°C. Lysates were then clarified with a spin. Clarified lysates were combined with 2 μL of AGO4 antibody (Agrisera) or 2uL of rabbit IgG as a mock IP (Cell Signaling Technology), and rotated at 4°C for 1 hour. Immune complexes were harvested with 40 μL of Protein G Dynabeads (Thermo), pre-washed in 1X TBS, rotating 30 min at 4°C. Beads and immune complexes were washed 3X with 1mL of cold wash buffer (50 mM Tris pH 8, 150 mM NaCl, 5 mM MgCl_2_, 0.5 mM DTT). Immunoprecipitated small RNA (bound by AGO4) was released from beads and isolated by TRIsure (Bioline) extraction. All RNA recovered from AGO4 IPs went directly into small RNA library preparation.

The small RNA library was prepared using TruSeq Small RNA Library Preparation kit (Illumina) as described above for total small RNAs with the exception of using 14 cycles of PCR amplification. Small RNAs were processed exactly as described earlier and accumulation was calculated in reads per million (RPM) sequenced reads for each size class of small RNAs in AGO4-IP and mock-IP samples. The small RNA enrichment was calculated as the ratio of RPM values for AGO4-IP over mock samples for each size class. This enrichment value is displayed as heatmap in Figure 2C and Supplemental Figure 6.

### Plasmid-Safe PCR Assay

Frozen inflorescence tissue was ground with liquid nitrogen, and total DNA was purified using the QIAGEN DNeasy Plant Mini Kit. 1 ug of resulting RNased DNA was digested using 1 ul Plasmid Safe ATP-dependent DNase (Lucigen) in a 50 ul reaction for 16 hours at 37°C. Digestion was completed by twice adding 1 ul additional DNase and 1 ul additional ATP followed by two hours of incubation at 37°C for a total of 20 hours digestion. DNase was inactivated at 70°C for 30 min. Digested and undigested DNA from each line was amplified using PCR primers for the EVD coding region (Supplemental Table 1).

### Western Blotting

Frozen inflorescence tissue was ground with liquid nitrogen and resuspended in lysis buffer (50 mM Tris pH 8, 150 mM NaCl, 5 mM MgCl_2_, 10% glycerol, 1% IGEPAL, 0.5 mM DTT, 1 mM PMSF, 1X Roche protease inhibitor cocktail) and homogenized with mixing for 15 minutes at 4°C. Lysates were clarified with a spin, combined with 2X loading buffer, denatured, and then loaded onto a 4%-20% gradient Tris-Glycine gel (Thermo). Protein was transferred from the gel to a PVDF membrane using the BioRad semi-dry transblot. Membranes were blocked for 1 hour at room temperature in 3% milk powder 1X PBS-T. Primary antibodies, which include Pol V (Wierzbicki laboratory), AGO4 (Agrisera), and ACT11 (Agrisera), were all diluted 1:1000 in 3% milk 1X PBS-T solution and incubated on blots overnight. Washes were performed at room temperature with 1X PBS-T. Anti-rabbit secondary antibody (Sigma) was used for visualization of Pol V and AGO4, while anti-mouse secondary (Sigma) was used for ACT11. Blots were visualized using HRP chemiluminescence (Thermo), with exposures ranging from 5 seconds to 5 minutes.

### Chromatin IP and Quantitative PCR

Nuclei were crosslinked as follows: frozen inflorescences (300 mg per biorep) were ground with liquid nitrogen and resuspend in nuclear isolation buffer (10 mM HEPES, 1 M sucrose, 5 mM KCl, 5 mM MgCl_2_, 0.6% Triton X-100, 0.4 mM PMSF, 1X Roche protease inhibitor cocktail) and homogenized with mixing for 15 minutes at 4°C. Methanol-free formaldehyde (Pierce) was added to a final concentration of 1% with end-over-end mixing for 15 minutes at room temperature. Formaldehyde crosslinking was quenched with glycine (125 mM), rotating 5 minutes at room temperature. Crosslinked nuclei were then filtered through two layers of Miracloth to remove large particles before centrifugation at 3000 x g for 15 minutes at 4°C. The resulting nuclear pellet was resuspended in wash buffer (10 mM Tris pH 8, 0.25 M sucrose, 10 mM MgCl_2_, 1 mM EDTA, 1% Triton X-100, 1X Roche protease inhibitor cocktail) and nuclei were cleaned and pelleted at 12,000 x g for 10 min at 4°C. The final clean nuclear pellet was resuspended in 1 mL nuclear lysis buffer (20 mM Tris pH 8, 2 mM EDTA, 0.1% SDS, 1 mM PMSF, 1X Roche protease inhibitor cocktail) and sonicated on the Covaris E220 (150W peak power, 20% duty factor, 200 cycles/burst, 6 minutes). Insoluble debris was removed from the sonicated soluble chromatin by centrifugation at 12,000 x g for 10 minutes at 4°C. 30 µL of 5M NaCl and 20 µL of 30% Triton X-100 were both added to 920 µL of sonicated chromatin in order to sequester SDS before immunoprecipitation. 3% input by volume was set aside for each sample, and the remaining volume of sonicated chromatin was divided evenly among IPs and IgG negative controls for overnight immunoprecipitation with respective antibodies or IgG.

Pol V (Lagrange lab antibody) and Ser5P Pol II (Abcam) ChIPs were performed with 2 µL antibody, whereas AGO4 (Agrisera) and H3K9me2 (Abcam) ChIPs used 5 µL antibody per IP. Each experiment used the same volume of rabbit IgG as a negative control (Cell Signaling Technology). Immune complexes were collected using 40 µL of washed protein A/G magnetic beads (Pierce), rotating at 4°C for 2 hours. After collecting immune complexes, beads were washed at 4°C using one rinse and two 5-minute washes of each of the following buffers: low salt (150 mM NaCl, 0.1% SDS, 1% Triton X-100, 2 mM EDTA, 20 mM Tris pH 8), high salt (500 mM NaCl, 0.1% SDS, 1% Triton X-100, 2 mM EDTA, 20 mM Tris pH 8), LiCl buffer (250 mM LiCl, 1% Igepal, 1% sodium deoxycholate, 1 mM EDTA, 10 mM Tris pH 8), and TE + 0.1% Igepal. Chromatin was eluted from the beads using 250 µL elution buffer (1% SDS, 0.1 M NaHCO_3_) at 65°C for 15 minutes with agitation. Overnight reverse crosslinking was accomplished with the addition of 20 µL 5M NaCl at 65°C for all samples including inputs. Proteinase K digestion was performed by the addition of 10 µL 0.5 M EDTA, 20 µL 1 M Tris pH 7, 20 ug Proteinase K (Thermo Scientific) and 50 ng RNase A, incubating at 42°C for one hour. DNA was then purified using DNA Clean and Concentrator-25 columns (Zymo Research) and eluted in 100 µL of elution buffer.

Quantitative PCR was performed using primers in Supplemental Table 1 and Sso Universal SYBR (BioRad). Percent input was calculated by first normalizing the Ct values of the diluted input samples to 100% input by the following calculation: Ct_100%_= Ct_diluted_ - (log(dilution factor, 2)). Then the IgG and IP Ct values were normalized to this new 100% input Ct value using the 2^-ΔΔCt^ method, which represents % input. For Pol V and AGO4 ChIP, the *pol V* mutant and *ago4/6/9* mutants (*ago4-4* allele, a complete null) were respectively used to calculate background levels for each of these antibodies. These background levels were averaged for each PCR target. They were then subtracted from the % input values of each of the other genotypes. Mean and standard deviations of the biological replicates are shown as bar graphs with error bars.

### ChIP-seq and RIP-seq Data Analysis

Raw reads were downloaded from NCBI GEO (GSE52041 and GSE70290), trimmed for adapters and mapped to the Araport11 Arabidopsis genome. The ChIP-seq reads were mapped using *Shortstack* (parameters: --nohp --mmap f --bowtie_m all). The RIP-seq reads were mapped using *Soapsplice 1*.*10*^53^ using parameters: -t 10300 (maximum distance between two segments) as mentioned in the original study^9^. For Figure 3A, reads were counted using *Bedtools* for each individual locus. To determine enrichment, the normalized ratio of IP over mock sample reads was calculated (for both ChIP-seq and RIP-seq). In eukaryotes, organelle-to-nucleus DNA transfer is known to occur which has resulted in some mitochondrial genes being duplicated and part of the nuclear genome. To avoid such genes causing an artefact of Pol V enrichment in the negative control dataset, we analyzed only those mitochondrial genes which do not have any similarity to nuclear genes.

In Figure 3C, genome-wide Pol V enriched loci were determined using the *Macs2* ChIP peak caller^54^. Pol V fold enrichment was calculated using the ratio of IP over mock samples. The two replicates were averaged for wt Col and *suvh2/suvh9* mutant. The loci that had ≥2-fold Pol V enrichment in wt Col and only background levels of Pol V enrichment in *suvh2/suvh9* were categorized as SUVH2/SUVH9-fully dependent Pol V loci. The loci that retained at least ≥2-fold Pol V enrichment in *suvh2/suvh9* mutants were categorized as SUVH2/SUVH9-partially dependent Pol V loci. In Figure 4C, AGO4 enrichment was calculated as a ratio of IP over mock normalized read accumulation. For genome-wide analyses, the two biological replicates were averaged, whereas the replicates are also displayed individually in Figure 3A.

## Data Availability

Raw Illumina sequencing data produced for this study is available from NCBI as GSEXXXXXX. Additional small RNA datasets were downloaded from GSE118705. Processed BSAS data of DNA methylation levels is available as Supplemental Dataset 1. Sanger sequencing results are available as Supplemental Dataset 2.

## Code Availability

The custom script used to generate RdDM Strength plots is available on Github - https://github.com/jpeasari/Dot-Plot-Anaysis-OpenCV.

## Material Availability

Biological materials can be obtained from the corresponding author without restriction.

## Acknowledgements

The authors thank Diego Cuerda-Gil, Seth Edwards, and Peng Liu for generating small RNA sequencing libraries. We thank Thierry Lagrange, Craig Pikaard and Andrzej Wierzbicki for sharing NRPE1 seed and antibody resources, Mike Axtell for *ago4-4/6/9* seed, and Yijun Qi for *dcl1/2/3/4* seed. We thank Jennifer Mele at The Ohio State Genomics Shared Resource, the University of Delaware DNA Sequencing & Genotyping Center and the Donald Danforth Plant Science Center Data Science Facility for computing support. We also thank the Donald Danforth Plant Science Center Plant Growth Facility, Gary Posey at the Center for Applied Plant Sciences Greenhouse, and Emily Yoders-Horn and David Snodgrass at the Biological Sciences Greenhouse. This work is supported by NSF grant MCB-1908521 to R.K.S.

## Author Contributions

Conceptualization - M.J.S., K.P., R.K.S., A.D.M.; Methodology - M.J.S., K.P., R.K.S., A.D.M.; Software - J.P., K.P.; Investigation - M.J.S., R.K., L.L.M., H.P., K.P., A.D.M.; Resources - A.Y.H.; Data Curation - K.P.; Writing - M.J.S., K.P., R.K.S. and A.D.M.; Funding Acquisition - R.K.S.

