## Supplemental Figures and Table for "An siRNA-guided Argonaute protein directs RNA Polymerase V to initiate DNA methylation"

#### Suppplemental Figure 1

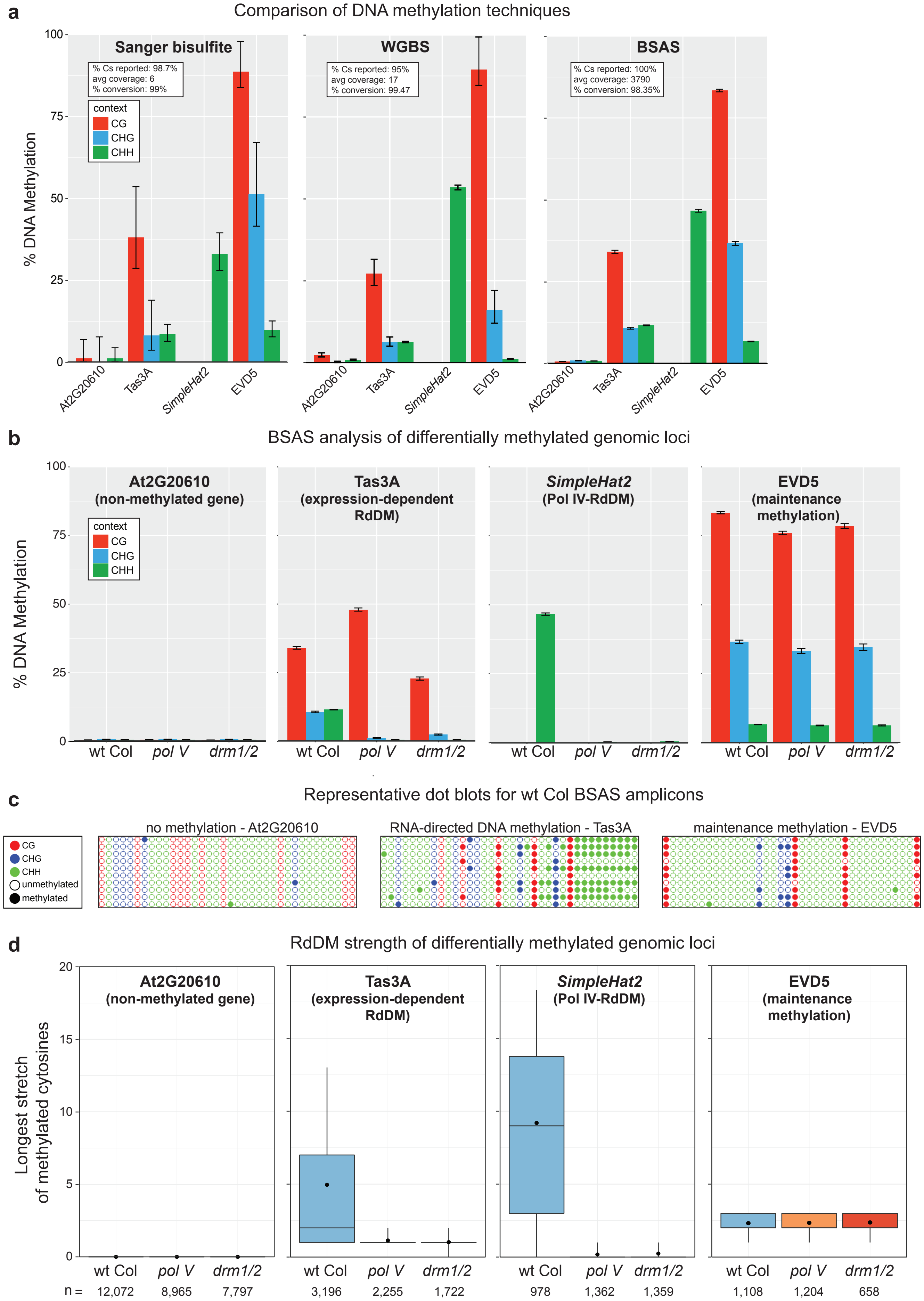

**Supplemental Figure 1 - BSAS and RdDM strength validation**

A. Comparison of three DNA methylation assays (Sanger sequencing of bisulfite converted DNA, whole genome bisulfite sequencing (WGBS), and bisulfite amplicon sequencing (BSAS)) at endogenous loci representing four different methylation mechanisms and scenarios: a non-methylated gene (At1G20610), an expression-dependent RdDM locus (Tas3A), a Pol IV-RdDM TE (*SimpleHat2*), and a maintenance methylation TE (the endogenous EVD5 - At5TE20395). The amplicons interrogated by Sanger sequencing are the same as in BSAS. Error bars represent 95% Wilson confidence intervals. BSAS results in much tighter confidence intervals than either Sanger sequencing or WGBS. The bisulfite conversion rate is empirically determined by assaying the methylation of the un-methylated genic locus At2G20610 for each technique. The *SimpleHat2* amplicon only has CHH methylation data, because it lacks CG and CHG cytosine contexts.
B. BSAS of four loci in wt Col, *pol V* and *drm1/2* plants confirms that BSAS is appropriate for assaying DNA methylation at all types of methylated loci from part A. Tas3A and *SimpleHat2* are targeted by RdDM and therefore are dependent on Pol V and DRM1/2 for DNA methylation, while the endogenous EVD5 element undergoes maintenance methylation and is therefore not dependent on these RdDM factors.
C. A snapshot of the dot blot analysis of DNA methylation data produced for 10 individual sequences using *Kismeth*[49]. Circles represent the cytosines in the DNA sequences, and filled circles are methylated cytosines. The circle color refers to the cytosine sequence context. The signature of RdDM is horizontal stretches of consecutively methylated cytosines as seen for Tas3A, but not for the maintenance methylation locus of endogenous EVD5.
D. Box plots of RdDM strength summarize the number and length of methylation stretches on individual sequenced reads. The endogenous EVD5 element that undergoes maintenance methylation (see part B-C) is an example of a region that has DNA methylation present but has low RdDM strength. N = the number of sequenced reads that contribute to the box plot data for each sample.

#### Supplemental Figure 2

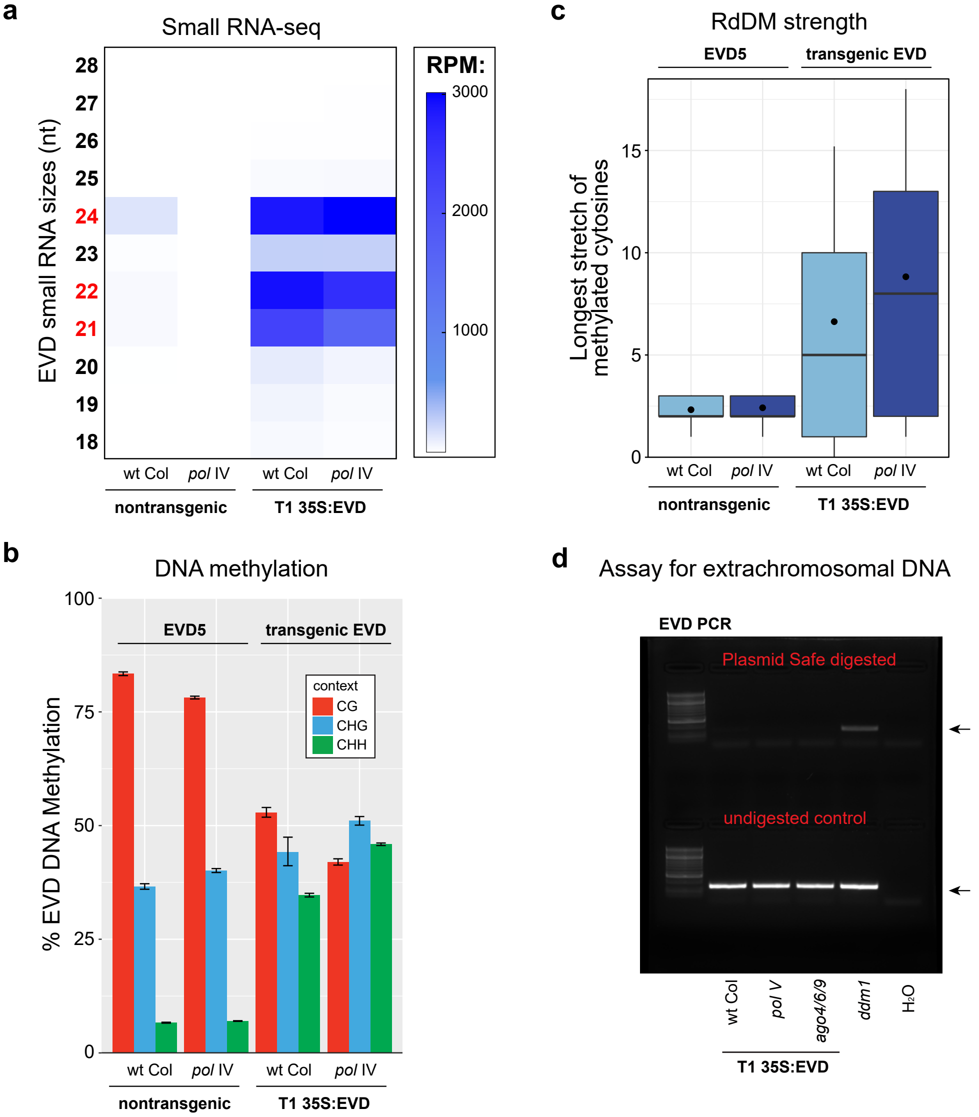


**Supplemental Figure 2 - The 35S:EVD transgene is not influenced by siRNAs generated by endogenous EVD elements and is not mobile**

A. EVD small RNA accumulation from non-transgenic wt Col and *pol IV* mutants, as well as both these lines with the 35S:EVD transgene. There is a low level of Pol IV-dependent EVD siRNAs in non-transgenic wt Col, while the siRNAs from 35S:EVD are not Pol IV-dependent.

B. BSAS of the endogenous EVD5 element and 35S:EVD. Pol IV is not responsible for the methylation of either locus.

C. Box plots of RdDM strength for the lines from B. This analysis demonstrates that 35S:EVD is undergoing expression-dependent initiation of DNA methylation only. It is not being targeted in trans from endogenous EVD via an identity-based mechanism of DNA methylation through Pol IV-dependent siRNAs. Therefore, 35S:EVD represents a transgene that is efficiently and reproducibly targeted in cis for de novo expression-dependent DNA methylation.

D. PCR amplification of Plasmid Safe DNase-digested DNA from 35S:EVD T1 lines (wt Col, *pol V,* and *ago4/6/9*) and a nontransgenic *ddm1* (F6) positive control. Undigested control DNA is shown in the bottom row. Plasmid Safe DNase digests genomic (linear) DNA leaving only circular, extrachromosomal DNA available for amplification (55). Only the *ddm1* positive control, in which endogenous EVD has been previously shown to transpose (56), has evidence of mobility. These results demonstrate that the 35S:EVD transgenic lines contain no extrachromosomal copies of either 35S:EVD or endogenous EVD. Arrows represent expected size of EVD amplicon. EVD primers amplify both endogenous EVD and 35S:EVD.

**Supplemental Figure 3**

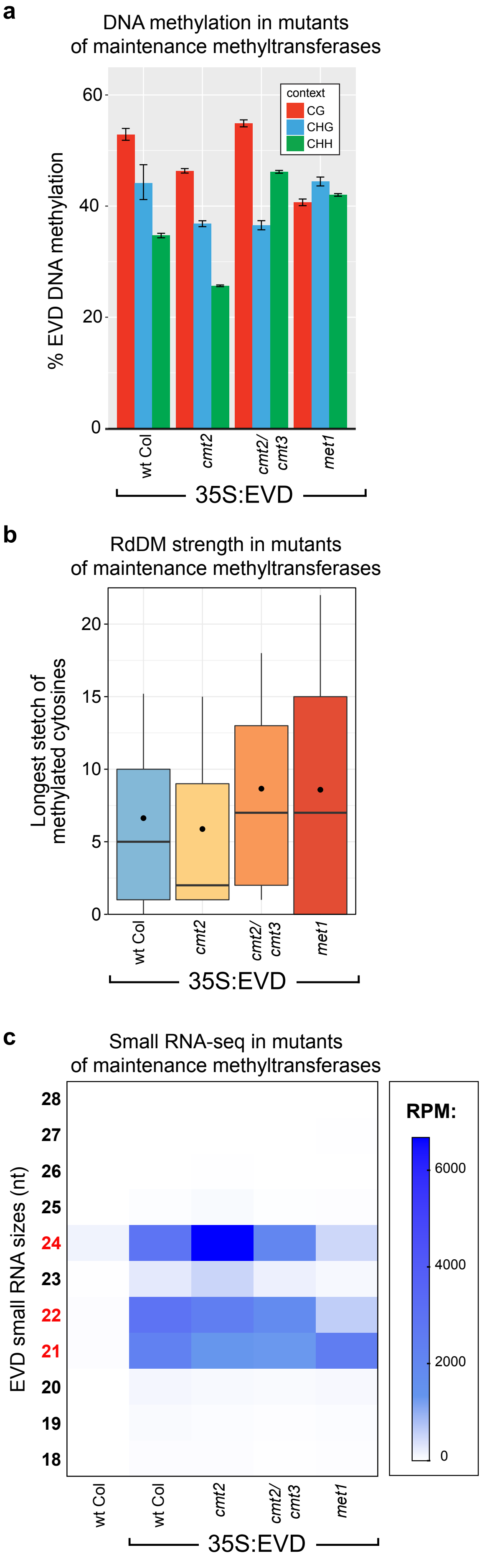


**Supplemental Figure 3 - Methylation of 35S:EVD is not dependent upon maintenance methylation factors**

A. BSAS of T1 35S:EVD in mutants of maintenance DNA methylation factors.

B. Box plots of RdDM strength in the same lines as part A.

C. EVD siRNA accumulation in the same lines as part A-B.

Supplemental Figure 4

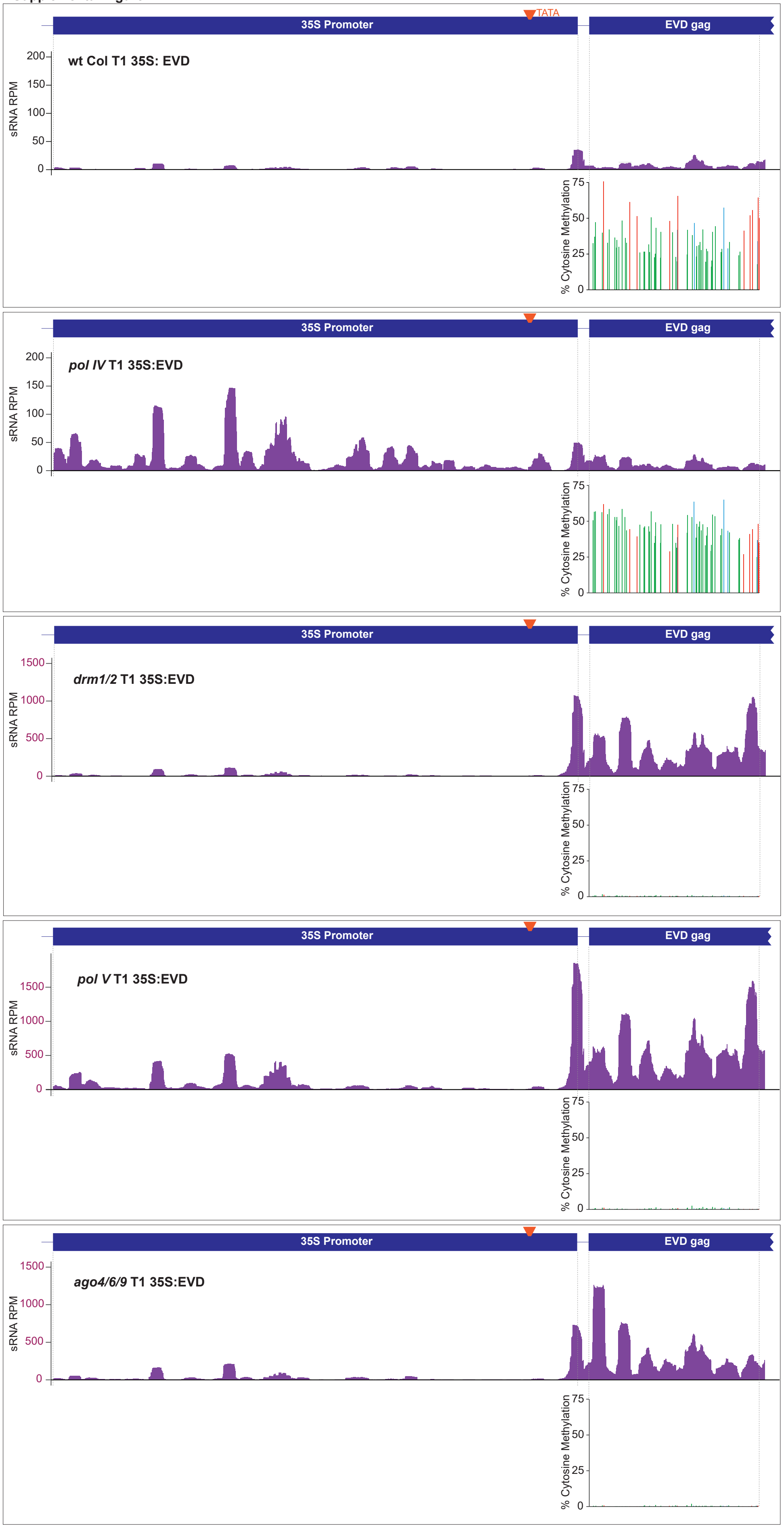


**Supplemental Figure 4 - Track images of DNA methylation and siRNA accumulation**

DNA methylation and siRNAs (combined 21, 22 and 24 nt) aligned to the 35S:EVD transgene. 35S promoter, TATA box, and EVD coding region are annotated. The siRNA alignments for T1 35S:EVD in *pol V, ago4/6/9*, and *drm1/2* have a larger Y-axis to accommodate for the increased siRNA production in these lines. BSAS methylation data from EVD amplicons is shown below each corresponding siRNA track and aligned to the primer set used for amplification. As shown, despite high levels of siRNA production in the T1 35S:EVD lines *pol V, ago4/6/9,* and *drm1/2*, there is no methylation for the assayed EVD region of the 35S:EVD transgene.

#### Supplemental Figure 5

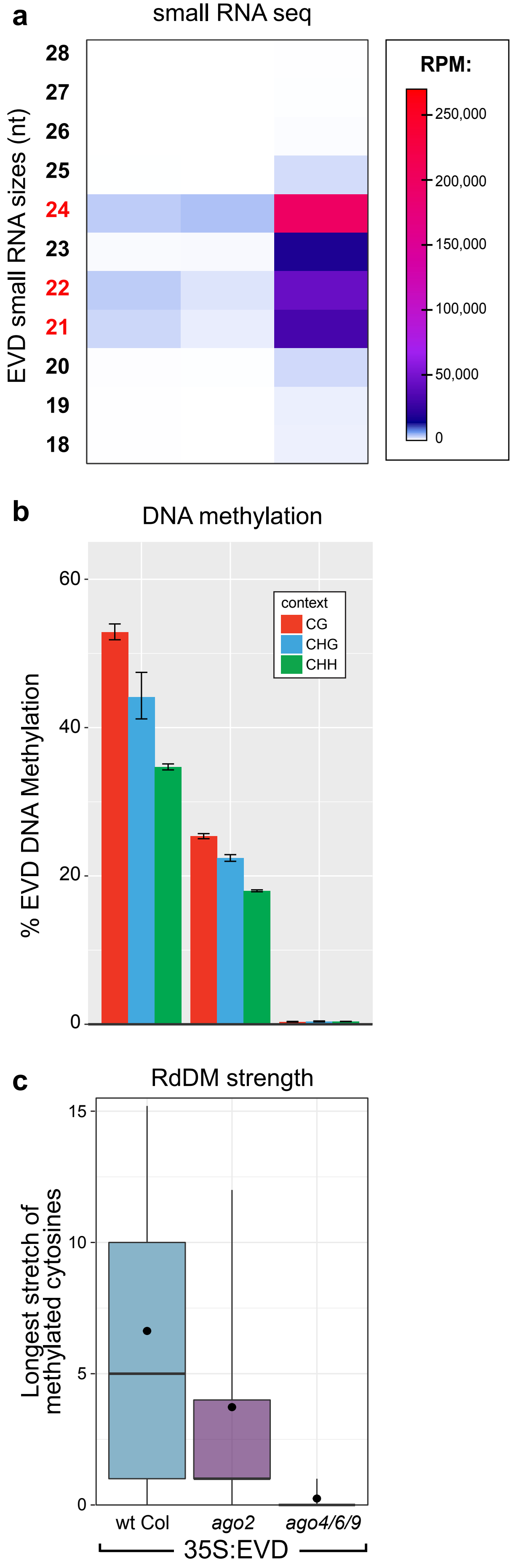


**Supplemental Figure 5 - Analysis of *ago2* mutant plants during the first round of RdDM**

A. EVD siRNA accumulation of the T1 35S:EVD transgene in *ago2* mutants and mutants of the ago4/6/9 line, which is shown for comparison.

B. BSAS of T1 35S:EVD in *ago2* mutants.

C. Box plots of RdDM strength in the same lines as part B.

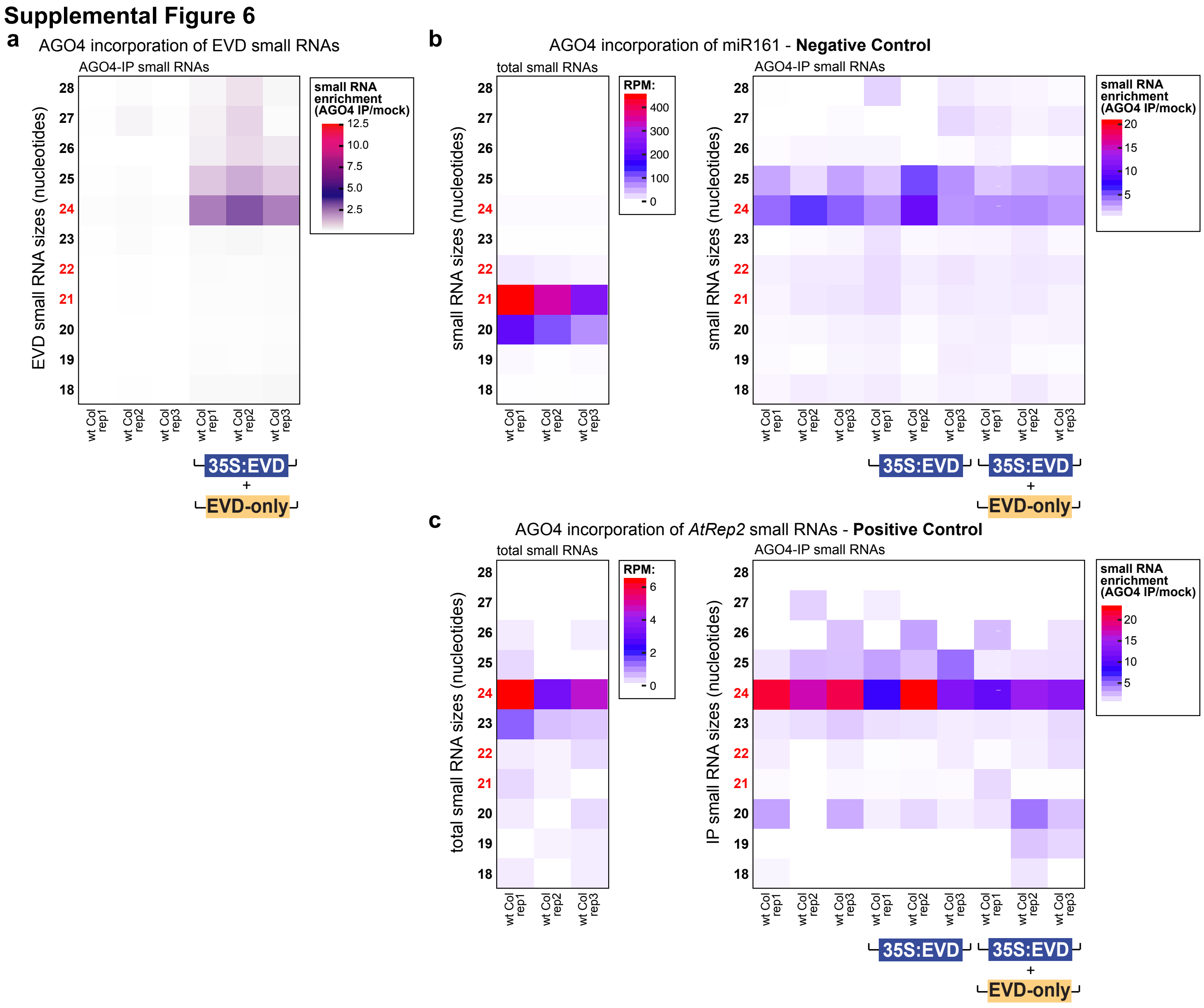


#### Supplemental Figure 6 - AGO4-IP small RNA sequencing

A. Heatmap of EVD small RNA enrichment in AGO4 for plants from Figure 6G, for plants containing both the 35S:EVD transgene and the no-promoter EVD-only transgene. Enrichment is calculated as the ratio of small RNA accumulation (RPM) in AGO4-IP over mock-IP samples for each size class of small RNA.

B. Negative control demonstrating the quality of AGO4 enrichment analysis for samples from Figure 2C and Supplemental Figure 6A. Heatmap of total small RNAs (RPM) is shown on the left whereas AGO4-IP small RNA enrichment is shown on the right side. Only small RNAs mapping to the miR161 region are displayed. miR161 accumulates in the cell as a 20-21 nt microRNAs (left), and this size class is not incorporated into AGO4 (right). Instead, low levels of a 24-25 nt version of miR161 are incorporated into AGO4.

C. Positive control demonstrating the quality of AGO4 enrichment analysis for samples from Figure 2C and Supplemental Figure 6A. The positive control *AtRep2* is a TE that generates 24 nt siRNAs (see total small RNAs heatmap on the left), which are incorporated into AGO4 (see AGO4-IP enrichment heatmap on the right).

### Supplemental Figure 7

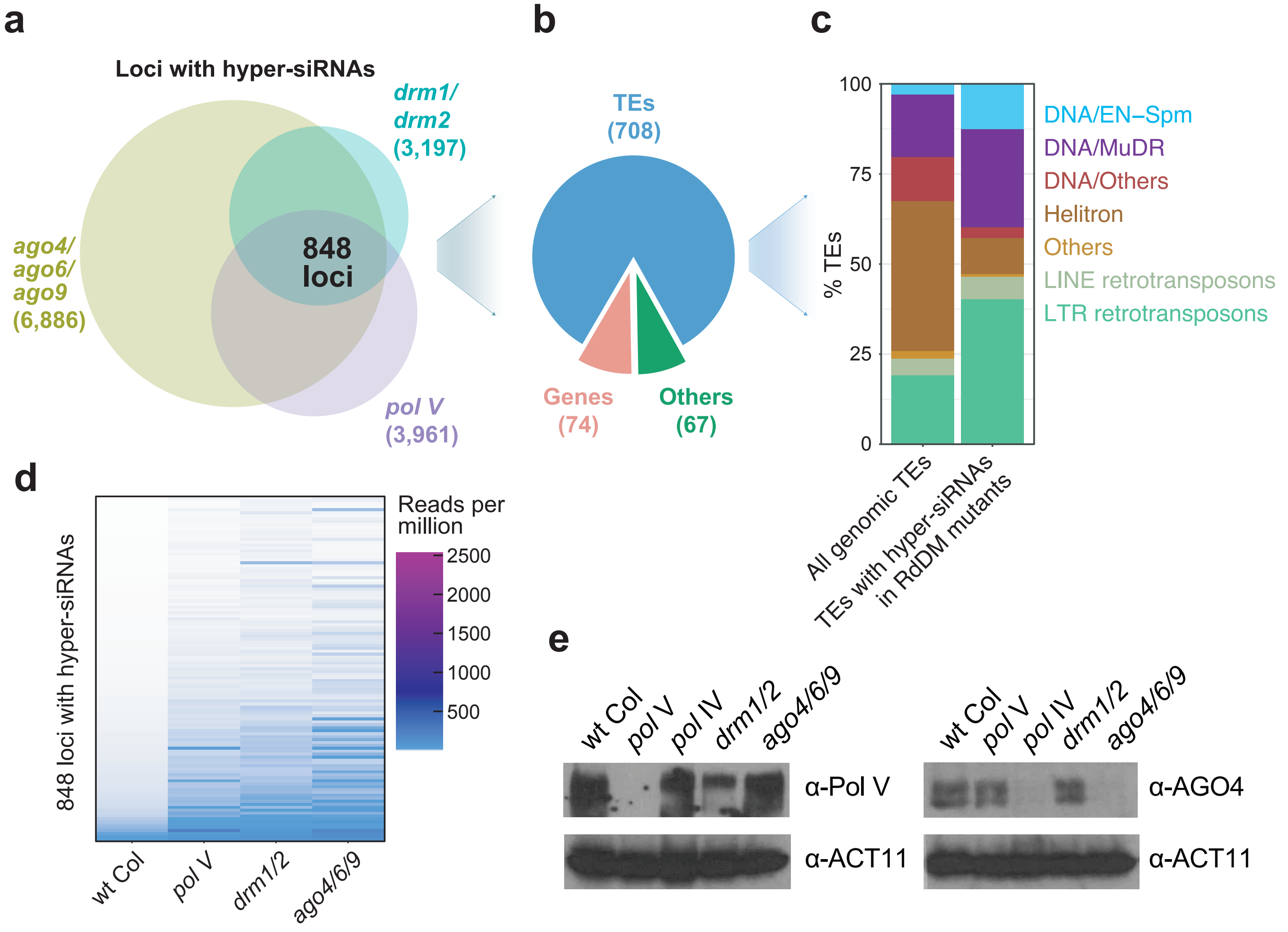


**Supplemental Figure 7 - siRNA and protein accumulation in mutants of downstream RdDM factors**

A. Venn diagram of loci with at least a 2-fold increase in siRNA production (combined 21, 22 and 24 nt) in *pol V, drm1/2* and *ago4/6/9* mutants. 848 loci have hyper-accumulation of siRNAs in all three mutant combinations.

B. Pie chart of the annotations of the 848 regions.

C. Distribution of TE superfamily annotations of the 708 TE regions form part B.

D. Heatmap of the siRNA accumulation (combined 21, 22 and 24 nt) in 848 loci with increased siRNAs from part A.

E. Western blots of Pol V and AGO4 protein accumulation in the mutant genotypes used in ChIP experiments. ACT11 protein accumulation is shown as a loading control.

**Supplemental Figure 8**

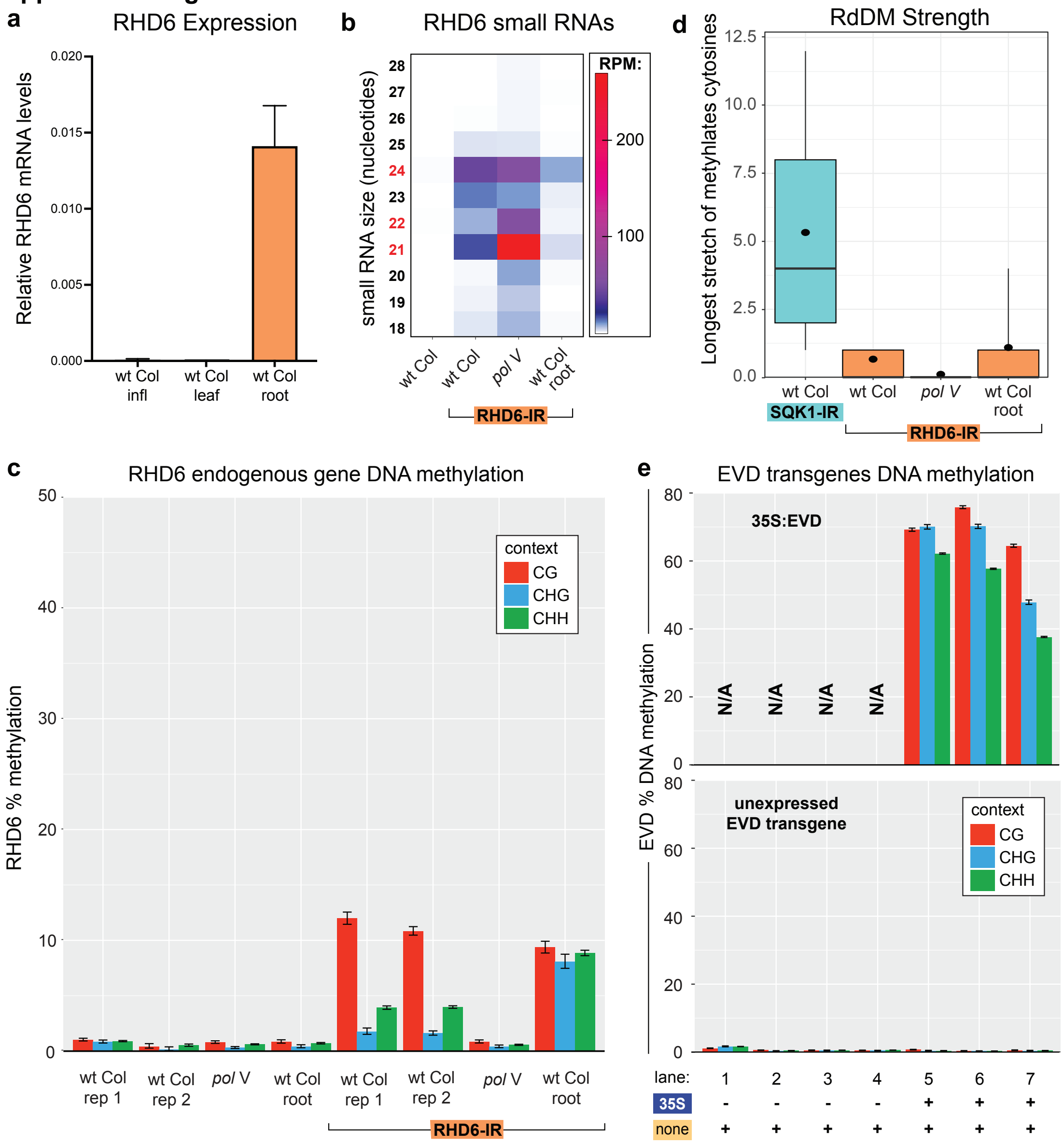


**Supplemental Figure 8 - Targeting RdDM to a dynamically expressed gene and unexpressed transgene**

A. mRNA accumulation of the ROOT HAIR DEFECTIVE 6 (RHD6, At1G66470) gene measured by qRT-PCR. The RHD6 gene is expressed only in the root hairs and developing embryo (57,58), and not in the inflorescence (Infl) tissue used in our other experiments. The error bar represents the standard deviation.

B. Heatmap of small RNA sequencing in T1 transformants. The endogenous RHD6 gene does not produce siRNAs, and upon the addition of the 'RHD6-IR' transgene in the T1 generation, 21-24 nt siRNAs accumulate from the IR transgene.

C. BSAS for T1 plants with the RHD6-IR. The IR siRNAs target a low level of methylation at the RHD6 endogenous gene. This methylation is dependent on Pol V, demonstrating that it underwent RdDM. The higher level of CG methylation compared to CHG and CHH in the inflorescence tissue suggests that this methylation is being maintained, rather than actively targeted as in the root sample that shows equal levels of methylation at all cytosine sequence contexts.

D. Box plots of RdDM strength for transgenic plants from C. The strength of RdDM is low in the inflorescence tissue analyzed compared to the SQK1-IR system. In inflorescence tissue, the low RdDM strength demonstrates that the unexpressed RHD6 is not actively undergoing RdDM in the inflorescence tissue. Rather this methylation is being maintained from RdDM earlier in development, likely in the embryo. Although the average is similar between inflorescence and root, the 90th percentile whisker shows that there are some sequenced reads from root undergoing RdDM at similar levels as the SQK1-IR, even with fewer IR-derived siRNAs accumulating in the root (part B).

E. Biological replicates of unexpressed EVD transgenes from Figure 6H. BSAS for the 35S:EVD transgene (top) and the second unexpressed EVD transgene (bottom) are from the same transgenic individuals. N/A = not applicable, as that transgene is not in this plant line.

#### Supplemental Table 1. Primer sequences and alleles used in this report

**Primers and sequences used in this report**

| Figure | Experiment | Target | Forward primer sequence | Reverse primer sequence |
|---|---|---|---|---|
| 1A | Cloning 35S:EVD | EVD5 - At5TE20395 | AGGCGGCCGCACTAGTATGGAGACTTCTCAAAAGATGATTAC | AGCTCAAGCTAAGCTTTCAAGAGTGAGATAGATCCACAAG |
| 1A/B, 2A/F, 6F/H, S2B/C, S3A/B, S4, S5B/C, S8E | BSAS | EVD coding region (35S:EVD) | GGAAGTTYATTTYATTTGGAGAGG | TAATCRCCTTCTTCACTTCAAACAC |
| 1A/B, 2A/F, 5C/D, 6C/F/H, S1A/B/C/D, S2B, S3A, S4, S5B/C, S8C/D/E | BSAS | control gene At2g20610 | GTTGYTGATTATATGAAYYGAGATYTT | TTAATTACAACCATARCCACARTRTTCTC |
| 2D/E, 3B, 4D/E | ChIP: H3K9me2, Pol II, Pol V, AGO4 | EVD coding region | CTATTCTAGTCGACCTGCAGGCG | AAGGACAGCTTGATCCTCTTGG |
| 2D | ChIP: Pol II | SimpleHat2 | AACTCAACCCATGAACCCTAATG | CCATTAACCATTACAACCCAACTC |
| 2D/E, 3B, 4D/E | ChIP: H3K9me2, Pol II, Pol V, AGO4 | ASX2 | AGGCAACCCAAACAATGAAGTG | GATCGTTGGACTCTCGATAGCTG |
| 2E | ChIP: H3K9me2 | *Athila6* TE | AGAAAGAAGAAGGCTGCACTCG | CACCGCTCTGATGATACTCGAC |
| 3B, 4D/E | ChIP: Pol V, AGO4 | positive control At5g52070 | CATCTGATTCTTAACACCACCTACTCA | ATGTCCTGAGCTGCCACGTT |
| 5A | Cloning SQK1-IR | Full sequence synthesized by Thermo Fisher | 35S Pro...CGCTCGAGCAGAGAGATCAA...SQK1...ACGAGCCCTTGGTAAGGAAATA…PDK<br>Intron...TTGGTCTAGAGATTTCGTCTAGATCGTTCA… SQK1...ACTAGTCCCTAGAGTCCTGCTT…OCS TERM | |
| 5C/D, 6A/C | BSAS | SQK1 amp 1 | TGTTGGGAAYTTAAAGGATTA | CTTTCTCARAARCARATTTCACCAC |
| 5A, S8 | Creating Hygro-IRs | 35S-HygroR-35ST | AATTTTCGCGTGAAACCTGTCGTGCCAGC | TAAAATGAGCTCTAACACATTGCGGACGTTTTT |
| 6A | BSAS | SQK1 amp 2 | AAAAYYAAAAGTAAAGTTTAATTAG | TTCCATTCCCRTCTATTAATTTTT |
| 6A/C | BSAS | *sqk1-1* deletion allele | AGAAGATGAAGATTYATTGGGTTTG | CTTTCTCARAARCARATTTCACCAC |
| 6B | qRT-PCR | SQK1 | TCACTGTGCTTAAGGGCGTT | TGCAATGTCAGGGGCGTAAG |
| 6B/E | qRT-PCR | control gene ASX2 | AGGCAACCCAAACAATGAAGTG | GATCGTTGGACTCTCGATAGCTG |
| 6A/B/C | Creating gRNAs for *sqk1-1* (4 primers) | SQK1 gRNAs amplified from pCBC-DT1T2 | ATATATGGTCTCGATTGATATGCGTTATCTGCCACGGTT<br>TGATATGCGTTATCTGCCACGGTTTTAGAGCTAGAAATAGC | AACTGCATATGTGGTAGAGACCCAATCTCTTAGTCGACTCTAC<br>ATTATTGGTCTCGAAACTGCATATGTGGTAGAGACCCAA |
| 6A/B/C | Cloning NapP:dsRED:NosT | Nap Promoter/Nos Terminator | CTTCATCGGTGATTGATTCCTTTAAAGACTTAT | GATCTAGTAACATAGATGACAC |
| 6A/B/C | Adding MfeI ends to NapP:dsRED:NosT | Nap Promoter/Nos Terminator | AAAAAACAATTGCTTCATCGGTGATTGA | AAAAAACAATTGGATCTAGTAACATAGATG |
| 6A/B/C | Genotyping for *sqk1-1* deletion | SQK1 promoter | GAGAAGCCAGTCCCAACAGG | GGGTCTCGGTCGTGAAGAAG |
| 6A/B/C | dCas9 Cloning from pDIRECT | dCas9 | ATGGATAAGAAGTACTCTATCGGACTCG | TCAAACCTTCCTCTTCTTCTTAGGATCAG |
| 6A/B/C | Creating gRNAs for dCas9 targeting (4 primers) | SQK1 (F) and RHD6 (R) gRNAs amplified from pCBC-DT1T2 | ATATATGGTCTCGATTGATACTCAAAAATTAATAGAGTT<br>TGATACTCAAAAATTAATAGAGTTTTAGAGCTAGAAATAGC | AACCCACCGGCATAAGCTGCATCAATCTCTTAGTCGACTCTAC<br>ATTATTGGTCTCGAAACCCACCGGCATAAGCTGCATCAA |
| 6A/B/C | In Fusion Primers for gRNAs + dCas9 | dCas9 into pMJS064 | CGCCGAATTAATTCGAGCTCCTAGGTTAATCACCACCGAGCTC | AAATTATCAGATCCGAGCTCGTAAAACGACGGCCAGTGC |
| 6A/B/C | In Fusion Primers for Rbsc Term | Rbsc Term into pMJS064 + dCas9 | TGGTGATTAACCTAGGAGCTTTCGTTCGTATCATCGG | GAATTAATTCGAGCTCCTAGCAATTGGCAAGTCATAAAATGCA |
| 6D | Cloning EVD-only | EVD5 - At5TE20395 | ATGATAATTCGAGCTCATGGAGACTTCTCAAAAGATGATTAC | AGCTCAAGCTAAGCTTTCAAGAGTGAGATAGATCCACAAG |
| 6D | Cloning T3A-EVD | T3A terminator | ATGATAATTCGAGCTCCAGGCCTCCCAGCTTTCG | CTTTTGAGAAGTCTCCATCTCGACACAAAAAGCCTATACTGTAC |
| 6D | Cloning T3A-EVD | EVD5 - At5TE20395 | ATGGAGACTTCTCAAAAGATGATTAC | AGCTCAAGCTAAGCTTTCAAGAGTGAGATAGATCCACAAG |
| 6E, S2D | qRT-PCR and Plastmid Safe PCR | EVD coding region | ACAAAAACGGTGTTGTGTGGTC | AAGGACAGCTTGATCCTCTTGG |
| 6F/H, S8E | BSAS | EVD coding region (EVD-only & T3A-EVD) | TAATGGTTTYTGAYGTATGTGYTTAGYT | TAATCRCCTTCTTCACTTCAAACAC |
| S1A/B/C/D | Sanger bisulfite sequencing & BSAS | Tas3A | GATTATTATAGTTGTAAAGAGTAA | CAACCATACATCAATAACAAACAA |
| S1A/B/D | Sanger bisulfite sequencing & BSAS | SimpleHat2 | AAAAAGTGGAATYYGTAAATATYTTAAGTTTGATG | TATTTTCATATTRTCAAAAAAACRTCTTCTTTCTC |
| S1A/B/C/D, S2B/C | Sanger bisulfite sequencing & BSAS | EVD5 | AGAGYATAAGATYATGTYTTTATG | TAATCRCCTTCTTCACTTCAAACAC |
| S8 | Cloning IR-RHD6 | Full sequence synthesized by Thermo Fisher | 35S Pro...CGCTCGAGATGTCAGAAGAAGCCGCC…RHD6...ACGAGCCCTTGGTAAGGAAA…PDK<br>Intron...TTGGTCTAGAGATTTCGTCTAGATCGTTTA…RHD6...CCCTACACGACTAGTCCCTAGAGTC…OCS Term | |
| S8A | qRT-PCR | control gene ACTIN 2 | CTCTTAACCCAAAGGCCAAC | ACACACCATCACCAGAATCC |
| S8A | qRT-PCR | RHD6 | CACGAGAGCTTTCCTCCTCC | TTCTGGCCTTGGGGAGTTTC |
| S8C/D | BSAS | RHD6 | GGAAGAGGAGGAATTTTGTTTTGA | CATTCATAACTTCRAARCTTCAAAC |

**Mutant alleles used in this report**

| Mutant | Allele | Background Ecotype | Figure |
|---|---|---|---|
| *pol V (nrpe1)* | SALK_017795 | Col | 1A/B, 2A/B, 3B, 4A/C/D, 5B/C/D, S1B/D, S2D, S4B/C/D, S7A/D/E, S8B/C/D |
| *suvh2/suvh9* | SALK_079574 / SALK_048033 | Col | 1A/B, 3C/D, 4E |
| *pol IV* | SALK_128428 | Col | 2A/B, S2A/B/C, S7E |
| *drm2* | SALK_150863 | Col | 2A/B |
| *drm1/drm2* | CS6366: 6530470652 / 6530470653 | WS | 1A/B, 2A/B/D/E, 3B, 4D, S1B/D, S7A/D/E |
| *ago4* | *ago4-6* (SALK_71772) | Col | 2A/B |
| *ago4* | *ago4-4* (FLAG_216G02) | WS | 2A/B |
| *ago6* | *ago6-2* (SALK_031553) | Col | 2A/B |
| *ago4/ago6* | CS66095: *ago4-1 / ago6-1* | mixed C24 and Ler | 2A |
| *ago4/ago6/ago9* | *ago4-4 / ago6-2 / ago9-1* | mixed Col and Ws | 2A/B/D/E, 3B/C/D, 4D/E, 5C/D, S2D, S5, S7E |
| *dcl1* | *dcl1-14* (SALK_056243) | Col | 2F/G |
| *dcl2/dcl4* | *dcl2-1* (SALK_064627) / *dcl4-2* (GABI_160G05) | Col | 2F/G |
| *dcl3* | *dcl3-1* (SALK_005512) | Col | 2F/G |
| *dcl2/dcl3/dcl4* | *dcl2-1 / dcl3-1 / dcl4-2* | Col | 2F/G |
| *dcl1/dcl2/dcl3/dcl4* | *dcl1-9/ dcl2-1 / dcl3-1 / dcl4-2* | Col | 2F/G |
| *cmt2* | SALK_012874 | Col | S3A/B/C |
| *cmt2/cmt3* | *cmt2-7 / cmt3-11* t | Col | S3A/B/C |
| *met1* | *ddm2-1* | Col | S3A/B/C |
| *ddm1* | *ddm1-2* | Col | S2D |
| *ago2* | SALK_003380 | Col | S5 |
| *sqk1-1* | Crispr promoter deletion allele, from this study | Col | 6A/B/C |
